# Helicase-driven unwinding defines the architecture of the leading-strand replisome

**DOI:** 10.64898/2026.05.20.726333

**Authors:** Megan C. DiIorio, Arkadiusz W. Kulczyk

## Abstract

DNA replication is a fundamental biological process that requires the coordinated activities of DNA helicase, DNA polymerase, and accessory proteins within the replisome. Although helicase-driven unwinding (HDU) has long been the accepted model for replisome progression, recent structural studies have proposed an alternative polymerase-driven unwinding (PDU) model, in which DNA polymerase physically separates duplex DNA. Here, we perform comprehensive structural and biochemical analyses to define the molecular architecture and functional interactions of the bacteriophage T7 leading-strand replisome to distinguish between these two mutually exclusive models. We show that the T7 DNA primase-helicase is positioned directly at the replication fork junction to drive DNA unwinding, whereas T7 DNA polymerase, in complex with its processivity factor *E. coli* thioredoxin, trails 11 to 15 nucleotides behind. Mutational analysis of T7 DNA polymerase reveals that a β-hairpin previously implicated as a strand-separation pin in PDU functions instead to stabilize DNA polymerase binding to the DNA template. Furthermore, we identify an essential interaction between an acidic amino acid patch in the exonuclease domain of T7 DNA polymerase and the primase domain of T7 DNA primase-helicase, which promotes highly processive leading-strand synthesis and is compatible only with the HDU model. Together, these findings provide evidence supporting helicase-driven unwinding and clarify the molecular organization of the T7 leading-strand replisome. Because the phage T7 replisome serves as a model for DNA replication, these results have broad mechanistic implications for replisomes across taxa.

**Significance Statement:** Processive DNA replication requires coordination between enzymes that unwind and copy DNA. Two mutually exclusive models have been proposed for replisome progression. In helicase-driven unwinding (HDU), DNA helicase separates the parental duplex at the replication fork to generate single-stranded templates for DNA synthesis. In contrast, an alternative model proposes polymerase-driven unwinding (PDU). Using structural and biochemical analyses, we investigated the architecture and functional interactions of the leading-strand replisome from bacteriophage T7. Our results show that DNA helicase performs DNA unwinding, while DNA polymerase follows to synthesize the nascent strand. We identify specific intermolecular interactions essential for leading-strand replication that are compatible only with the HDU model. These findings support helicase-driven unwinding and provide insights with broad mechanistic implications across taxa.

## Introduction

DNA replication is a fundamental biological function essential for the inheritance of genetic information in all living organisms. This highly accurate and efficient process depends on the assembly and coordination of a large, multiprotein complex known as the replisome. Replication occurs at a replication fork, where DNA helicase unwinds the double-stranded DNA (dsDNA) to generate single-stranded DNA (ssDNA) templates for copying by DNA polymerases. Due to the reverse polarity and antiparallel orientation of DNA, the leading-strand DNA is synthesized continuously, while the lagging-strand is replicated discontinuously through the formation of short Okazaki fragments. Physical interactions between replication proteins coordinate the simultaneous synthesis of both strands.

Due to its simplicity, the bacteriophage T7 replisome is an attractive model system that has been studied extensively (1). The T7 replisome can be reconstituted with only four proteins that capture all fundamental activities of more advanced replication systems: gene protein 5 DNA polymerase (gp5) and its processivity factor *E. coli* thioredoxin (trx), bifunctional gene protein 4 DNA primase-helicase (gp4), and gene protein 2.5 single-stranded DNA-binding protein (gp2.5). Interactions between these four proteins are sufficient to orchestrate DNA replication, whereas more complex systems rely on many additional factors. The human replisome, for example, requires over 30 proteins to replicate the genome (2).

The intrinsic dynamics and complexity of the replisome pose a challenge for structure determination (3–5). To date, no structure of a replication complex from any biological system has visualized all of its core protein components together with a complete trajectory of fork-shaped DNA. However, recent advances in cryo-electron microscopy (cryo-EM) (6–15) have enabled structural insights into the architecture of the T7 leading-strand replisome containing hexameric gp4, gp5/trx, and partially visualized DNA used for its assembly (3, 4). Upon analysis of the currently available cryo-EM maps, two mutually exclusive models emerge for the leading-strand replisome, with different implications for DNA unwinding, the directionality of replisome progression along DNA, and the mechanisms coordinating leading-strand synthesis. After integrating these structural data (3, 4) with biochemical (16–21) and single-molecule studies (22–27), we define the models as the helicase-driven unwinding (HDU) model (**Fig. 1A**) and the polymerase-driven unwinding (PDU) model (**Fig. 1B**). In the HDU model, supported by the first cryo-EM map of the bacteriophage T7 replisome from Kulczyk et al. (3), DNA primase-helicase migrates ahead of DNA polymerase and physically separates the duplex DNA at the replication fork junction. In the alternative PDU model supported by the cryo-EM maps from Gao et al. (4), DNA polymerase acts as the physical prow for dsDNA unwinding. The HDU model has been the prevailing view of replisome progression in all systems for decades; nevertheless, new structural data challenges this perspective.

**Fig. 1.**
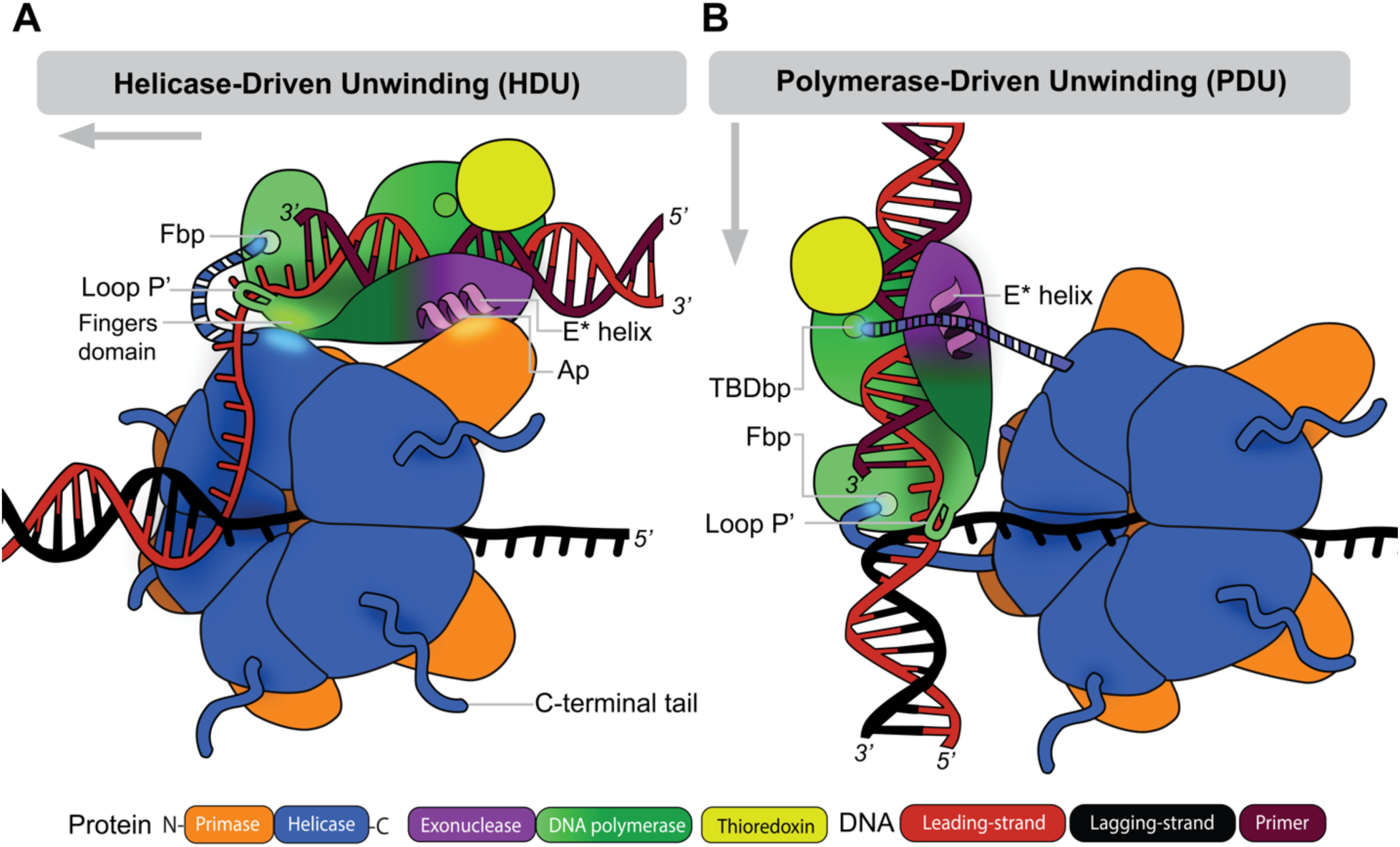
Conflicting models of the leading-strand replisome depict different modes for DNA unwinding. (**A**) In the helicase-driven unwinding (HDU) model of leading-strand replication, T7 DNA primase-helicase (gp4) rides ahead of T7 DNA polymerase (gp5/trx) to unwind dsDNA. Gp5/trx is bound to the backside of the gp4 hexamer relative to the direction of unwinding. Both proteins perform concerted unwinding and synthesis along the long axis of DNA. This protein geometry implies gp4 unwinds dsDNA at the fork junction, and gp5/trx engages the newly exposed leading-strand ∼10 nucleotides (nt) upstream. The complex is stabilized by multiple contacts between gp5/trx and gp4. A patch of acidic residues (Ap) in α-helix E* of the gp5 exonuclease domain interacts with the primase domain of gp4, while the gp5 fingers domain contacts the adjacent helicase subunit of gp4. In addition, the C-terminal tail of gp4 may remain in contact with the front basic patch (Fbp) of gp5 to maintain processive synthesis. (**B**) In the polymerase-driven unwinding (PDU) model, DNA polymerase is directly involved in dsDNA unwinding, while gp4 plays an accessory role. The P’ loop of gp5/trx acts as a separation pin to unwind the parental duplex, which requires a 2-nt ssDNA gap between the fork junction and primer-template. Gp4 does not directly interact with dsDNA and translocates along the unwound lagging-strand perpendicular to the axis of DNA unwinding. Coordination of leading-strand synthesis is mediated through interactions between the C-terminal tail of gp4 and Fbp of gp5/trx. Additional contacts may also be maintained between another C-terminal tail and the basic exchange patch of gp5/trx (TBDbp). Gray arrows indicate the direction of DNA unwinding.

Here, we integrated biochemical studies with detailed structural analysis to differentiate between the two models by systematically investigating their unique features and functional implications, namely: (i) the relative positioning of DNA primase-helicase and DNA polymerase within the functional leading-strand replisome assembled on forked-shaped DNA, (ii) the trajectories of the forked-shaped DNA through the complex, (iii) the protein-protein and protein-DNA interactions, and (iv) the enzymatic activities required for DNA unwinding and synthesis during coordinated leading-strand replication.

## Results

### Structural Analysis of the HDU and PDU Models

Although multiple cryo-EM maps corresponding to phage T7 leading-strand complexes, representing HDU (EMDB ID: 8565) and PDU (EMDB IDs: 0391-0395), have been deposited to the EMDB (**SI Appendix**, **Fig. S1** and **Movie S1**), these depositions are not accompanied by atomic coordinates submitted to the RCSB PDB. Likewise, atomic coordinates corresponding to the entire bacteriophage T7 replisome bound to DNA resembling a replication fork are not available (3, 4), highlighting the need for further biochemical and structural analysis to elucidate the principles underlying the mechanism of coordinated DNA synthesis. Thus, to facilitate the interpretation of protein and DNA densities in the HDU (3) and PDU (4) maps, we performed rigid-body docking of atomic models using a combination of UCSF ChimeraX (28), Phenix2.1 (29), and Coot1.1 (30) (see **Materials and Methods**). Briefly, atomic coordinates of DNA polymerase (PDB ID: 1T7P for HDU; PDB ID: 6N7W for PDU) and DNA primase-helicase (PDB IDs: 1CR0, 1NUI for HDU; PDB IDs: 6N7V,1NUI for PDU) were first docked into the maps using UCSF ChimeraX (28). These docked models were then iteratively refined using Phenix2.1 (29) and Coot 1.1 (30).

Analysis of the resultant models highlights key structural features that mediate the coordination of leading-strand synthesis according to the HDU and PDU models (**Fig. 2 A** and **B**, respectively). In the HDU model, gp4 is positioned to unwind the parental duplex (**Fig. 1A**) and provide the ssDNA template for the leading-strand gp5/trx (3). The leading-strand replisome advances along the long axis of the parental duplex, with gp5/trx bound to the backside of the gp4 hexamer relative to the direction of unwinding. This geometry implies a 10-15 nucleotides (nt) long gap between the fork junction and the leading-strand primer bound by gp5/trx.

**Fig. 2.**
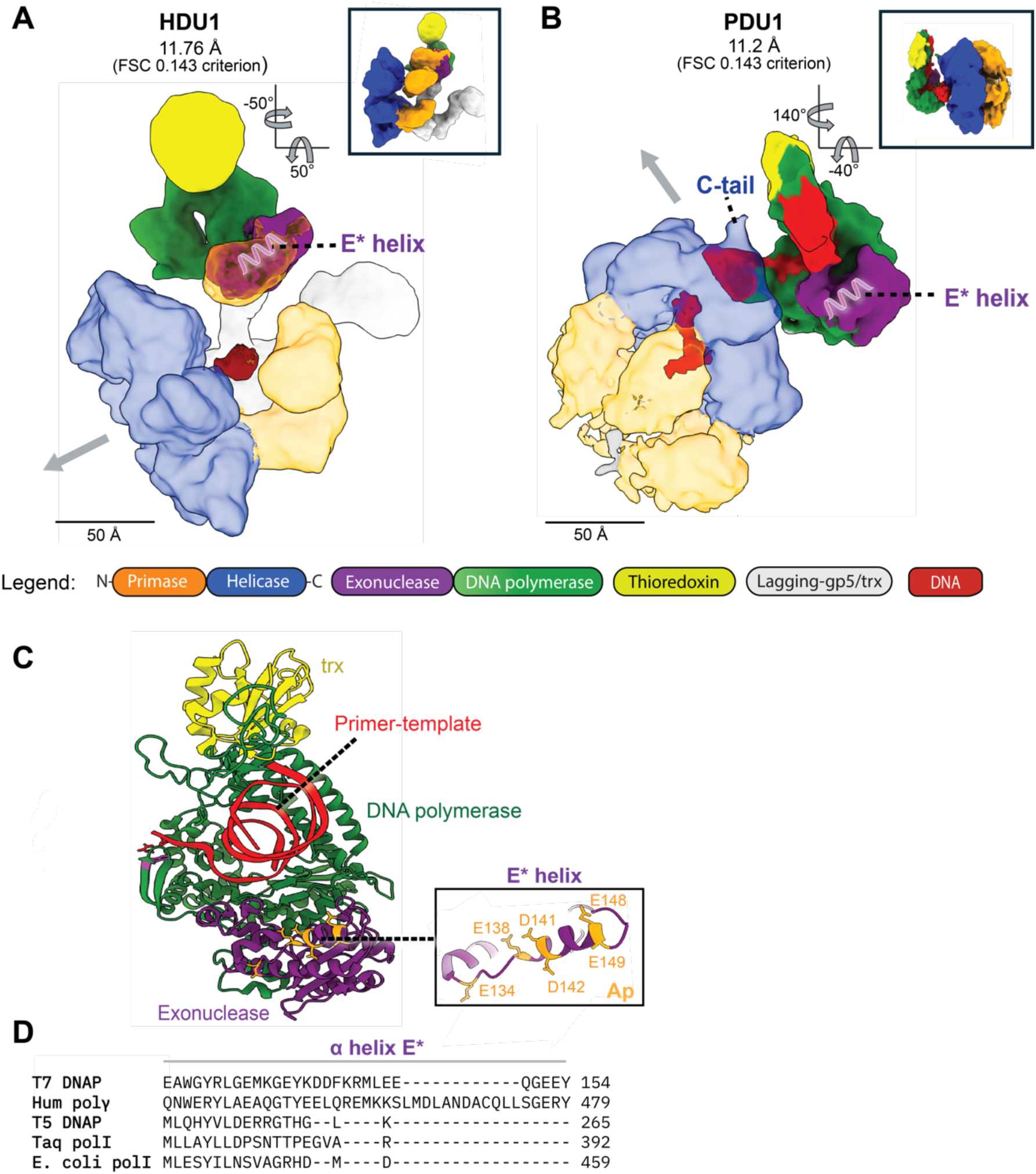
Structural comparison of HDU and PDU cryo-EM maps of the T7 leading-strand replisome reveals different positions of gp5/trx and the E* helix. (**A**) In the HDU-supporting cryo-EM map (EMD-8565), the E* helix (shown as a magenta cartoon) of gp5 contacts the primase domain of gp4 (orange), (3). For clarity, densities representing helicase (blue), primase (orange) and lagging-strand gp5/trx (gray) are shown as transparent. (**B**) In the map representing the PDU (EMD-0391), gp5/trx is located on the C-terminal face of the gp4 hexamer, and the E* helix is in close proximity to the parental duplex (4). Gray arrows indicate the directions of dsDNA unwinding. (**C**) Crystal structure of the gp5/trx-primer-template complex (PDB ID: 2AJQ), (52). The inset shows the E* helix, with residues belonging to the acidic patch (Ap) shown in orange. (**D**) Amino acid sequence alignment of selected DNA polymerases showing the unique E* helix (Glu126-Gln150). Included polymerases: T7 DNA polymerase (T7 DNAP), Homo sapiens DNA polymerase γ (Hum polγ), T5 DNA polymerase (T5 DNAP), Thermus aquaticus DNA polymerase I (Taq polI), and Escherichia coli DNA polymerase I (E. coli polI). Sequences were aligned using Clustal Omega 1.83 (86).

Importantly, the 11.7 Å (FSC 0.143 criterion) cryo-EM structure from Kulczyk et al. (3) depicting the HDU model shows that gp5/trx makes extensive contacts with both the primase and helicase domains of adjacent gp4 subunits through its exonuclease and fingers domains, respectively (**Fig. 2A**). Notably, the former interaction is mediated through the α-helix E* (residues Glu126 - Gln150), (**Fig. 2C**) in the exonuclease domain of DNA polymerase (3), (**Fig. 2A**). The E* helix is unique to gp5/trx among Pol I family polymerases (**Fig. 2D**), and it contains a patch of solvent-exposed acidic residues (Glu134, Glu138, Asp141, Asp142, Glu148, and Glu149) we refer to as the acidic patch (Ap). Although not visualized in the cryo-EM structure, supporting biochemical and single-molecule studies indicate that another interaction between the acidic C-terminal tail of gp4 and a solvent-exposed front basic patch (Fbp) of gp5/trx may be maintained in HDU (25, 27, 31), (**Fig. 1A**).

In contrast, the PDU model positions gp5/trx directly at the fork junction, where it is proposed to unwind dsDNA (**Fig. 1B**). In this model, gp4 translocates along ssDNA perpendicular to the axis of DNA unwinding without directly engaging the parental duplex. This geometry produces an approximately 90° bend in the lagging-strand before entry into the central helicase channel, placing gp4 in a noncanonical orientation at the replication fork and suggesting a limited role in duplex unwinding. Instead, gp4 is proposed to stabilize the unwound lagging-strand, while gp5/trx simultaneously performs DNA unwinding and leading-strand synthesis (4, 19). For gp5/trx to unwind the parental duplex, a strictly defined gap of 2 nt between the fork junction and the leading-strand primer is required (4, 19), (**Fig. 1B**). This arrangement is proposed to position an extended β-hairpin of gp5/trx as an intercalculating wedge at the fork junction, although not readily visualized in the cryo-EM maps determined by Gao et al. (4) at 9.6-13.8 Å resolution (FSC 0.143 criterion), (**SI Appendix**, **Fig. S2 A** and **B**). This element, which we define as the P′ loop (residues Glu575-Gln585), connects α-helix P to β-strand 10 in gp5 (32). The 2-nt gap between gp5/trx and the replication fork junction further implies that gp4 does not directly interact with the dsDNA. In contrast to the HDU model, leading-strand complex structures show that gp4 and gp5/trx in the PDU interact solely through electrostatic contacts between the flexible acidic C-terminal tails of gp4 and the Fbp and/or a second basic patch located in the thioredoxin-binding domain of gp5 (TBDbp). Interactions are observed at the Fbp in all five deposited maps in the EMDB, whereas contacts with the TBDbp are only observed in one of five maps (PDU1 in **SI Appendix**, **Fig. S1** and **Movie S1**), (4).

Importantly, no Coulomb maps of the HDU or PDU models visualized the full path of the DNA fork (**SI Appendix**, **Fig. S1** and **Movie S1**), (3, 4). Interestingly, the DNA ligands used in both studies were nearly identical, consisting of a stable tetraloop formed in the middle of the sequence to generate a fork-shaped DNA substrate originally designed by Kulczyk et al. to facilitate crystallization (33), differing only in the gap length between the fork junction and the primer-template (2 nt in the PDU and 10 nt in the HDU), (**Fig. 1**). Analysis of the PDU cryo-EM maps revealed Coulomb densities representing short fragments of the DNA fork (**Materials and Methods** and **SI Appendix**, **Fig. S1 A** and **B**). In all five PDU maps, density likely corresponding to ∼9 base pairs (bp) of parental dsDNA were resolved extending from the fork junction along the duplex, although the central tetraloop was not visible (**SI Appendix**, **Fig. S1B**). In four of five maps (PDU1-PDU4 in **SI Appendix**, **Fig. S1 A** and **B**), density likely representing fragments of the ssDNA lagging-strand was observed in the central helicase channel. However, it could not be traced continuously to the fork junction in any map, and no comparable density was present in PDU5. In addition, density corresponding to the dsDNA leading-strand primer-template bound by gp5/trx was present in all maps, although the 2-nt ssDNA gap between the fork junction and primer-template could not be unambiguously resolved (**SI Appendix**, **Fig. S1A**). As none of these features correspond to the density of a continuous DNA fork, it is not possible to determine whether the observed densities in the PDU maps represent a single DNA ligand or multiple ligands bound simultaneously, for example, gp4 bound to the lagging-strand of one ligand and gp5/trx binds the primer-template of a separate ligand, while the complex is stabilized only by protein-protein interactions.

The Coulomb map of the HDU complex, calculated from data collected using a CCD camera rather than a direct electron detector, which was not readily available at the time (34), also did not visualize the fork-shaped DNA. However, the hexameric organization of gp4, direct quantum dot labeling of the fork-shaped DNA in the cryo-EM samples, the presence of non-protein density within the helicase ring attributed to ssDNA (**SI Appendix**, **Fig. S1C,D** and **Movie S1**), and supporting biochemical evidence collectively supported the presence of DNA in the complex (3).

The absence of continuous density representing forked DNA in the cryo-EM maps, and the resulting inability to track its complete trajectory, prevents direct interpretation of DNA spacing, one of the crucial features distinguishing the HDU and PDU models, and highlights the need for further biochemical characterization of the DNA ligand. To address this, we designed DNA fork ligands and assessed their binding to gp4 and gp5/trx, providing a foundation for evaluating the HDU and PDU models.

### Design of a DNA Ligand Resembling the Replication Fork

A DNA scaffold is required for replisome assembly. A simplified replication fork for assembling a leading-strand replisome contains the following elements: (1) parental dsDNA, (2) a dsDNA leading-strand primer-template, (3) a ssDNA gap between the primer-template and the parental dsDNA, and (4) a ssDNA lagging-strand (**SI Appendix**, **Fig. S3**). Because protein-protein interactions within the replisome are transient, the architecture of the replication fork defines the three-dimensional arrangement of the proteins and the geometry of their interactions. Thus, careful design of the DNA substrate is required to reflect the physiological complex responsible for coordinated DNA synthesis. To define the DNA structural features required for optimal gp4 binding and evaluate how the DNA fork architecture supports gp4 positioning predicted by the HDU and PDU models, we examined the binding affinity of gp4 to DNA forks with varying lagging-strand 5′-overhang lengths (21-48 nt), 3′-overhang lengths (0-29 nt), and leading-strand ssDNA gap sizes (0-25 nt). In an analogous manner, we assessed the binding affinity of DNA polymerase to DNA forks with varying leading-strand ssDNA gaps (0-25 nt) and 5′-overhang lengths (0-20 nt). Finally, to define the spacing on the leading-strand that supports the simultaneous binding of DNA primase-helicase and DNA polymerase and the formation of specific protein-protein and protein-DNA interactions required for leading-strand synthesis, we examined DNA synthesis on DNA forks containing leading-strand gaps of 2-25 nt.

### Characterization of the DNA Primase-Helicase Binding Site on a DNA Fork

In the presence of deoxythymidine triphosphate (dTTP), gp4 binds to ssDNA as a hexamer and translocates in the 5’ to 3’ direction. We used the electrophoretic mobility shift assay (EMSA) to define the positioning of gp4 on DNA fork ligands with varying 5’-overhang lengths (21-48 nt) (**Fig. 3A**). DNA substrates were assembled by annealing a 5′-³²P-labeled 54-nt leading-strand oligonucleotide with lagging-strand oligonucleotides of varying lengths (42-69 nt), producing DNA forks with 21-to 48-nt 5’-overhangs. The gp4 used in these studies is a genetically altered protein (gp4-M4) with one amino acid substitution in the C-terminal helicase domain (E343Q) that eliminates gp4-M4 hydrolysis activity and consequently its ability to translocate along ssDNA. An additional three amino acid substitutions (P16K, D18H, N19R) were introduced in the N-terminal Zinc-Binding Domain (ZBD). Gp4-M4 displays approximately 5-fold higher binding affinity for ssDNA than wild-type gp4 (WTgp4), (35). In this reaction, DNA forks were incubated with increasing amounts of gp4-M4 in the presence of 0.1 mM dTTP at 37 °C for 10 minutes and analyzed by EMSA (**Fig. 3B-E**). A 21-nt 5’ overhang is suboptimal with the dissociation constant (K_d_) of 0.35 ± 0.01 µM (**Fig. 3B**, **3F-G**). Consistent with Kulczyk et al. (36), the data indicate the optimal 5’-overhang length for gp4-M4 binding is 27 nt, reflected by the K_d_ of 0.12 ± 0.01 µM (**Fig. 3C**, **3F-G**).The 33-nt and 47-nt overhangs are not optimal, as gp4-M4 binds these ligands approximately 3-to 4-fold less tightly compared to the 27-nt overhang (**Fig. 3D-G**). Given that the footprint of gp4 spans 25-30 nt (37), the 27-nt 5′-overhang, unlike the 33-nt and 48-nt 5′-overhangs, likely positions gp4-M4 in close proximity to the fork junction while providing structural features that stabilize gp4-M4 DNA binding. In contrast, the lower K_d_ measured with a 21-nt 5′-overhang indicates that ssDNA threading through the central channel of gp4 also contributes to DNA binding. Four high-molecular-weight (HMW) bands observed in the EMSA gels in **Fig. 3** represent protein-DNA complexes that likely correspond to distinct loading conformations, as reported previously (38). Interestingly, in the presence of the 27-nt and 33-nt 5′-overhangs, a single dominant HMW band appears, suggesting stabilization of the protein-DNA interaction near the fork junction (**Fig. 3 C** and **D**). These results are consistent with DNA footprinting experiments indicating that the helicase domain of gp4 is positioned close to the fork junction (18).

**Fig. 3.**
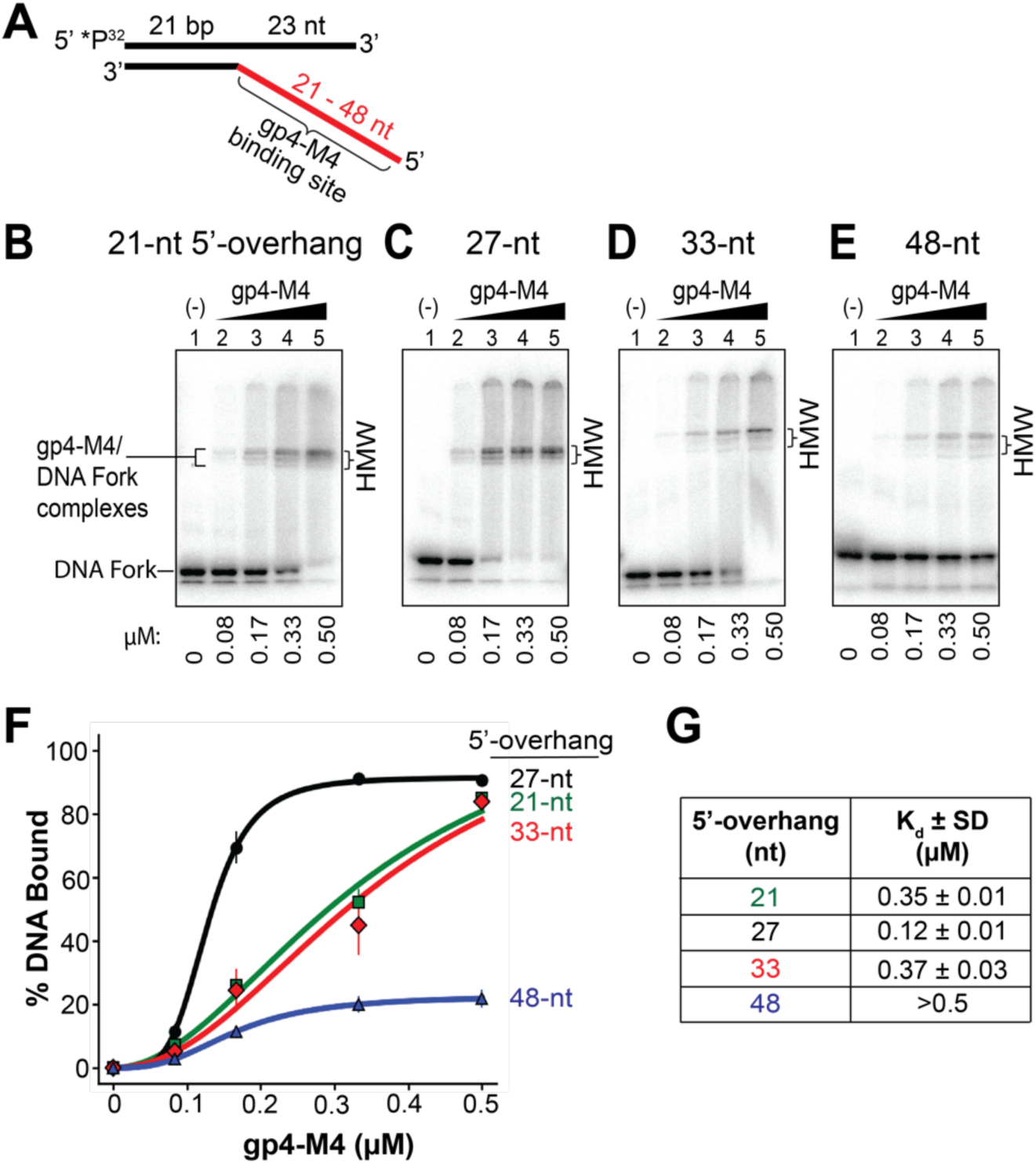
Characterization of DNA binding affinity of gp4-M4 for DNA forks with varying ssDNA 5’-overhangs. (**A**) DNA forks with a 21-bp duplex region and different 5′-overhang lengths (21-48 nt) were prepared by annealing a 54-nt leading-strand with a ^32^P-5’-end label to lagging-strands of varying lengths (42-69 nt). DNA forks (0.3 µM) were incubated with increasing concentrations of gp4-M4 in the presence of 0.1 mM dTTP at 37 °C for 10 minutes. (**B-E**) Representative EMSA gels revealing minimal protein stacking in the well. High-molecular-weight (HMW) bands representing gp4-M4-DNA complexes are indicated. Multiple bands likely represent different loading conformations that have been observed previously (38). Faint bands migrating below the unbound DNA fork on the gel represent excess unannealed ³²P-labeled leading-strand. (**F**) Quantification of DNA binding. The fraction of gp4-M4 bound to DNA was quantified by measuring the intensity of all HMW bands relative to total DNA in each lane and plotted against protein concentration. Error bars represent standard deviation of three independent experiments. (**G**) Table listing dissociation constants (K_d_).

We next assessed gp4-M4 binding to DNA forks with a 27-nt 5’-overhang and various ssDNA 3’-overhangs of up to 29 nt (**SI Appendix**, **Fig. S4**) as described above. Although, we find that gp4-M4 binds to DNA forks with similar affinities regardless of 3’-overhang length (K_d_ of 0.12 ± 0.01 µM for no overhang, 0.12 ± 0.02 µM for 3-nt overhang, 0.13 ± 0.06 µM for 10-nt overhang, and 0.16 ± 0.01 µM for 29-nt overhang), the maximal binding capacity (B_max_) is higher with longer overhangs (B_max_ of 73.57 ± 1.64 % for no overhang, 69.38 ± 1.20 % for 3-nt overhang, 81.75 ± 1.52 % for 10-nt overhang, and 88.29 ± 0.28 for 29-nt overhang), (**SI Appendix**, **Fig. S4G**). This observation suggests that the 3’-overhang may actively enhance protein accessibility to the binding site, possibly by facilitating more efficient loading.

Alternatively, it could influence the gp4-M4 off-rate kinetics. Interestingly, kinetics studies indicate gp4 interactions with the 3’-overhang are critical for unwinding activity (39–42), and gp4 cannot unwind a DNA substrate with 3’ blunt end (42). In support of our results, Jeong et al. (43) showed gp4 requires a 3’-overhang of at least 10 nt, and optimally 15 nt, to catalyze dsDNA unwinding. This length is consistent with distance between the primer-template and the replication fork junction required in the HDU model (**Fig. 1A**).

The HDU and PDU models predict different sized ssDNA gaps of approximately 10-15 nt (3) and 2 nt (4, 19), between the fork junction and the leading-strand primer-template, respectively (**Fig. 1**). Thus, defining this gap in the context of simultaneous binding of gp4-M4 and gp5/trx to the DNA fork is important for distinguishing between the two models. First, we examined gp4-M4 binding to DNA forks with various ssDNA gaps ranging from no gap to a 25-nt gap (**Fig. 4A-C** and **SI Appendix**, **Fig. S5A-F**). DNA fork substrates were assembled by annealing leading-strand oligonucleotides of varying lengths (41-66 nt) to a 48-nt lagging-strand oligonucleotide and a 20-nt primer labeled at the 5′ end with ³²P. Gp4-M4 binds weakly to DNA forks with no gap or a 1-nt gap, as demonstrated by a K_d_ of 0.83 ± 0.04 µM and > 0.83 µM, respectively, and reduced maximal binding (**Fig. 4 B** and **C**). Binding affinities of gp4-M4 to DNA forks increase when gap size increases with K_d_ of 0.52 ± 0.01 µM for the 3-nt gap and 0.51

**Fig. 4.**
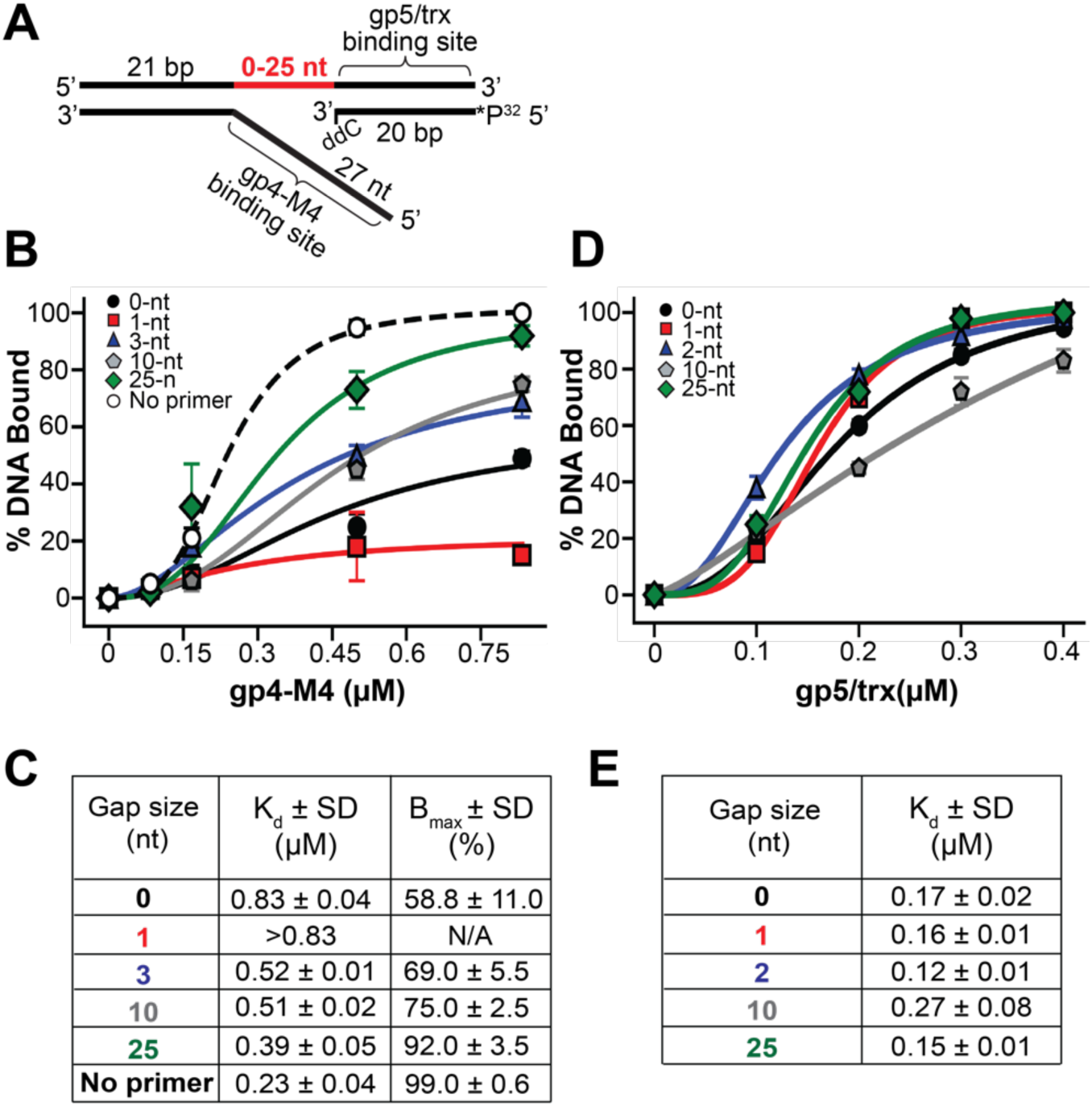
Characterization of DNA binding affinity of gp4-M4 and gp5/trx for DNA forks with varying ssDNA gap sizes. (**A**) DNA fork substrates containing ssDNA gaps of 0-25 nt (red) between the fork junction and the primer-template were assembled by annealing a 48-nt lagging-strand, a 20-nt primer (5′-end labeled with ³²P), and leading-strands of varying lengths (41-66 nt). To prevent primer extension, the 3′ end of the primer contains a dideoxycytidine (ddC). DNA ligands (0.3 µM) were incubated with increasing concentrations of gp4-M4 (0.08-0.83 µM, hexameric concentration) or gp5/trx (0.1-0.4 µM) in the presence of 0.1 mM dTTP at 37 °C for 10 min. (**B**) Fraction of DNA bound plotted as a function of gp4-M4 concentration. Error bars represent the standard deviation of three independent experiments. (**C**) Binding parameters for gp4-M4 derived from curve fits, including maximum binding (B_max_) and dissociation constants (K_d_). (**D**) Fraction of DNA bound plotted as a function of gp5/trx concentration. (**E**) K_d_ values for gp5/trx.

± 0.02 µM for the 10-nt gap (**Fig. 4 B** and **C**). Gp4-M4 exhibits the highest binding affinity among tested conditions for the DNA fork with a 25-nt gap (K_d_ of 0.39 ± 0.05 µM), (**Fig. 4 B** and **C**). This data reveals that the presence of a primer-template at 10 nt or less from the fork junction interferes with gp4-M4 binding, as reflected by an approximately 2-fold decrease in binding affinities and 1.5-to 2-fold decrease in B_max_ (**Fig. 4C**).

Taken together, these results suggest that at the replication fork, gp4-M4 is positioned close to the fork junction, and the primer-template, which provides a binding site for leading-strand gp5/trx, must be located more than 10 nt away to provide optimal gp4-M4 binding. This scenario is consistent with the HDU model of replication, in which the DNA helicase is positioned at the fork junction to directly unwind dsDNA while a trailing DNA polymerase extends the upstream primer-template. It is inconsistent with the PDU model, in which the opposite is true, namely the DNA polymerase binds at the fork junction, whereas the DNA helicase, bound to non-template ssDNA, trails behind gp5/trx as it extends the double-stranded primer-template (**Fig. 1**).

### Characterization of DNA Polymerase Binding Site on a DNA Fork

We next measured gp5/trx binding affinity to DNA forks with various gap lengths (0-25 nt) between the fork junction and primer-template (**Fig. 4 D** and **E**, and **SI Appendix**, **Fig. S5G-K**). We used a gp5 variant that contains two amino acid substitutions in the exonuclease domain (D5A and D7A). These mutations reduce exonuclease activity by a factor of 10⁶ and prevent degradation of DNA forks (32). For simplicity, we refer to this variant as gp5. DNA fork substrates were assembled as described in the previous section (**Fig. 4A**). A chain-terminating 2’,3’-dideoxycytosine triphosphate (ddCTP) was incorporated at the 3′ end of the leading-strand primer to prevent primer extension. Furthermore, the next incoming nucleotide specified by the template strand, dTTP, was included in the reaction mixture to force gp5/trx into the closed conformation, reflecting nucleotide incorporation during primer synthesis (32). We found that gp5/trx exhibits similar binding affinity across all tested DNA forks, regardless of gap size and distance from the fork junction (K_d_ of 0.17 ± 0.02 µM for no gap, 0.16 ± 0.01 µM for 1-nt gap, 0.12 ± 0.01 µM for 2-nt gap, 0.27 ± 0.08 µM for 10-nt gap, and 0.15 ± 0.01 µM for 25-nt gap), suggesting that structural features at the fork junction do not influence protein binding (**Fig. 4 D** and **E**). Although the K_d_ measured for the DNA fork with a 10-nt gap appears approximately 2-fold lower than K_d_ values determined for other gap sizes ranging from 0-2 nt, this measurement is associated with a relatively high standard deviation (**Fig. 4E**). Moreover, the K_d_ measured for a 25-nt gap is similar to those from the 0-2 nt range (**Fig. 4E**).

It has been speculated that in a PDU model of leading-strand synthesis, DNA polymerase facilitates dsDNA unwinding presumably through activity analogous to strand-displacement (4). If so, gp5/trx may directly interact with the displaced lagging-strand, analogous to the requirement of DNA primase-helicase for a 10-to 15-nt 3′-overhang representing the leading-strand for optimal DNA unwinding activity (43). Thus, we examined the strand-displacement activity of gp5/trx using DNA forks with varying 5’-overhang lengths (0-20 nt), and a 4-nt poly-A gap between the 3’ primer end and fork junction (**Fig. 5A**). The 4-nt gap was selected to facilitate separation of the reaction products on a sequencing gel, as longer DNA fragments would make this analysis more challenging. All deoxynucleotide triphosphates (dNTPs) were included in the mixture, except 2’,3’-dideoxyadenosine triphosphate (ddATP) was used in place of deoxyadenosine triphosphate (dATP) to stop strand-displacement by gp5/trx at the first dT in the template strand, located one nucleotide downstream from the fork junction (N_+1_; **Fig. 5A**).

**Fig. 5.**
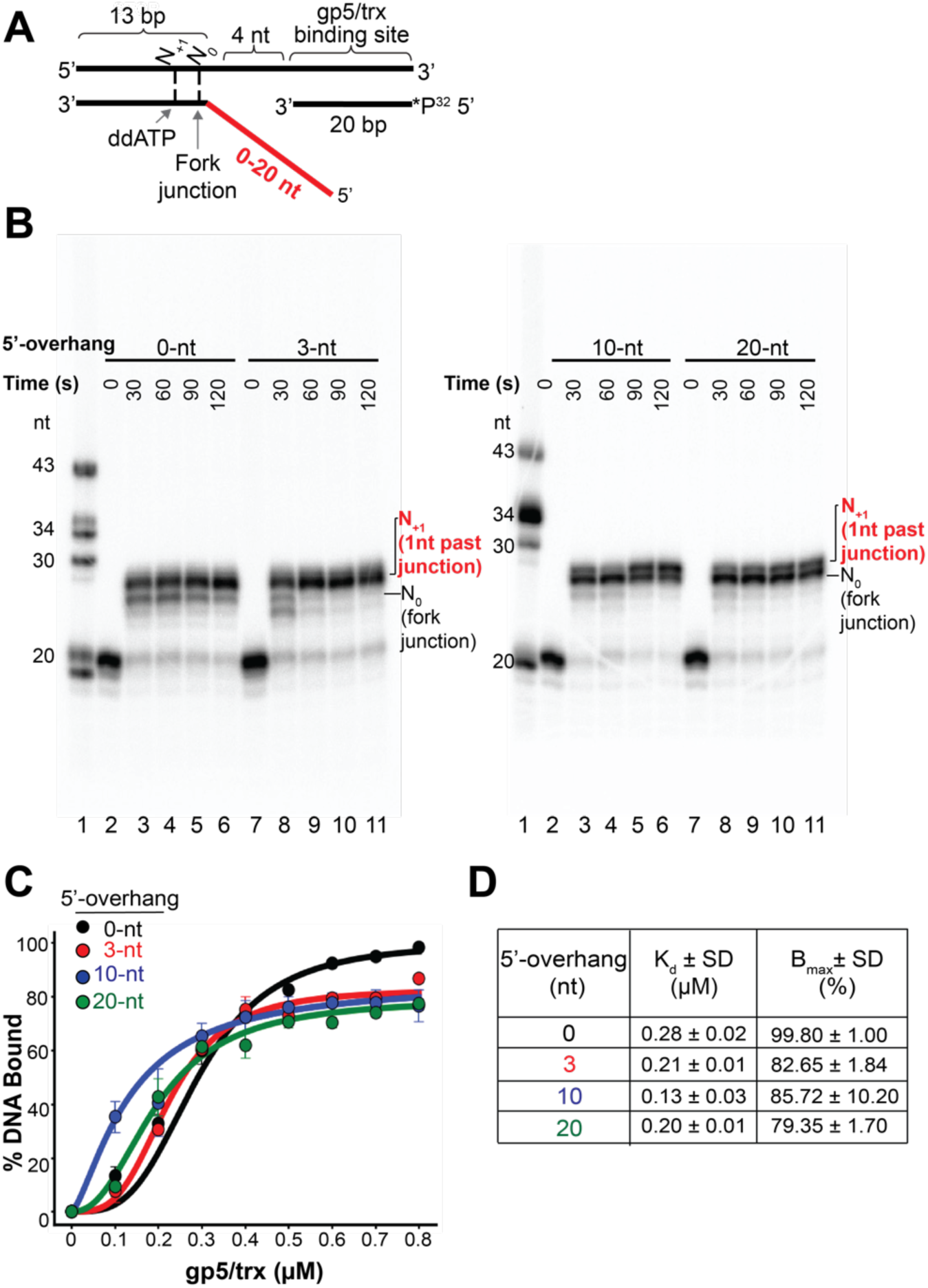
Strand displacement activity of gp5/trx is unaffected by 5’-overhang length. (**A**) DNA fork substrates containing different 5’-overhang lengths (0-20 nt) were obtained by annealing a 37-nt leading-strand, a 20-nt primer (5′-end labeled with ³²P), and lagging-strands of varying lengths (13-33 nt). Reactions were initiated by mixing equimolar amounts of gp5/trx and DNA with 0.1 mM of each dNTP, except dideoxyadenosine triphosphate (ddATP) was used in place of dATP to stop strand-displacement activity at position N_+1_ in the parental duplex. Reactions were incubated at 37 °C for the indicated time points (30, 60, 90, 120 s) and quenched with 20 mM EDTA. (**B**) The products were resolved on 8M, 20% polyacrylamide gels. Independent of 5’ overhang length, strand displacement synthesis stopped at the N_0_ or N_+1_ position, corresponding to 25- or 26-nt products. (**C**) EMSA analysis of gp5/trx stalled at the N_+1_ position on DNA forks with varying 5′-overhang lengths. Strand-displacement was performed under the conditions described in A, with reactions incubated for 2 minutes at increasing gp5/trx concentrations (0.1-0.8 µM). The fraction of DNA bound was quantified and plotted against gp5/trx concentration. Error bars represent standard deviation of three independent trials. (**D**) Table of dissociation constants (K_d_) and maximum binding (B_max_) derived from quantification.

This traps gp5/trx in a position reflecting its proposed DNA unwinding activity (4, 19). Reactions were carried out for up to 2 minutes, stopped at indicated time points with 20 mM EDTA, and the resulting DNA products were separated on denaturing polyacrylamide gels (**Fig. 5B**). Independent of the 5′-overhang length, all strand-displacement reactions terminated at the N_+1_ position (**Fig. 5B**, lanes 3-6 and 8-11). Notably, a substantial fraction of the reaction products corresponded to the N_0_ position, with the relative amounts of N_0_ and N_+1_ products varying as a function of time (**Fig. 5B**). At the N_0_ position, gp5trx incorporates the final nucleotide into the primer at the fork junction, nevertheless suggesting that its P′ loop is located such that it would participate in in dsDNA strand separation (**Fig. 1**). This result indicates that the length of the 5’-overhang does not influence the proposed dsDNA unwinding activity of gp5/trx at the measured timescale.

We next used EMSA to perform a similar experiment in which we evaluated gp5/trx complexes stalled on DNA forks as a function of the gp5/trx concentration (**SI Appendix**, **Fig. S6**), rather than time (**Fig. 5**). In this experiment, strand-displacement reactions were carried out for 2 minutes at varying gp5/trx concentrations in the presence of 5’-overhangs ranging from 0 to 20 nt. Reactions were resolved on native polyacrylamide gels, and the resulting band intensities were quantified and expressed as the percentage of shifted DNA as a function of protein concentration (**Fig. 5C**). Gp5/trx exhibits similar binding affinity across all tested DNA forks, regardless of the 5’-overhang length (K_d_ of 0.28 ± 0.02 µM for no 5’-overhang, 0.21 ± 0.01 µM for 3-nt, 0.13 ± 0.03 µM for 10-nt, and 0.2 ± 0.01 µM for 20-nt 5’-overhang). Although the K_d_ measured for the DNA fork with a 10-nt 5’-overhang (**SI Appendix**, **Fig. S6C**) is approximately 50% lower than K_d_ values determined for other 5’-overhang sizes ranging from 0-3 nt (**SI Appendix**, **Fig. S6A, B**), the K_d_ measured for a 20-nt 5’-overhang is similar to those from the 0-3 nt range (**SI Appendix**, **Fig. S6D**). Moreover, the highest B_max_ is observed for the DNA fork ligand without a 5’-overhang (**Fig. 5D**). This suggests that the presence of a 5’-overhang reduces accessibility of gp5/trx to its binding site. In summary, our results suggest that the structural features of DNA proposed to facilitate DNA polymerase-mediated unwinding, such as proximity to the fork junction and the presence and length of the displaced 5’ lagging-strand, either do not contribute to or may hinder DNA binding, respectively. These findings are inconsistent with the role proposed for DNA polymerase in the PDU model.

### The Stop-Trap Experiment Reveals that a 11-to 14-nt Gap on the Leading-strand is Required to Form a Functional Interaction Between DNA Polymerase and DNA Primase-Helicase

Coordinated leading-strand synthesis requires the formation of a stable replication complex, maintained through specific protein-protein and protein-DNA interactions that position the leading-strand replisome at a defined location relative to the fork junction. In **Fig. 4**, we showed that, unlike the efficient gp5/trx binding to DNA forks with varying gap sizes (0-25 nt, **Fig. 4 D** and **E**), the presence of a primer-template located ≤10 nt from the fork junction significantly interferes with gp4-M4 binding to the replication fork (**Fig. 4 B** and **C**). This observation is inconsistent with the PDU model, which requires a 2-nt gap (4, 19), but is consistent with the HDU model, which requires a 10-to 15-nt gap (3), (**Fig. 1**).

To further distinguish between these two models, we assembled gp5/trx and gp4-M4 on replication forks containing varying gap sizes (2, 4, 10, and 25 nt) and performed strand-displacement assays (**Fig. 6**). In these experiments, gp4-M4 stalled on the lagging-strand acts as a stop-trap for DNA polymerase and is expected to influence leading-strand synthesis through specific protein-protein and protein-DNA interactions. As in the previous section (**Fig. 5**), we have used ddATP to halt strand-displacement synthesis 2 bp into the parental duplex (position N_+1_). Alone, gp5/trx extends nearly all primers past the fork junction to position N_+1_ within the first 30 seconds of the reaction, regardless of gap size (**Fig. 6B-E**, lanes 3-7). However, when gp4-M4 was pre-bound to DNA forks containing 2-to 10-nt gaps, synthesis decreased significantly; after five minutes, extension to the N_+1_ position was reduced approximately 4-fold, 2.5-fold, and 5-fold for the 2-nt, 4-nt, and 10-nt gaps, respectively (**Fig. 6B-D**, lanes 8-12; summarized in **Fig. 6F**). Interestingly, when the gap length was 25 nt, gp5/trx did not extend any primers past the fork junction to the N_+1_ position in the presence of gp4-M4. Instead, DNA synthesis was halted after incorporation of 11 to 14 nt into the extended primer (**Fig. 6E**, lanes 8-12).

**Fig. 6.**
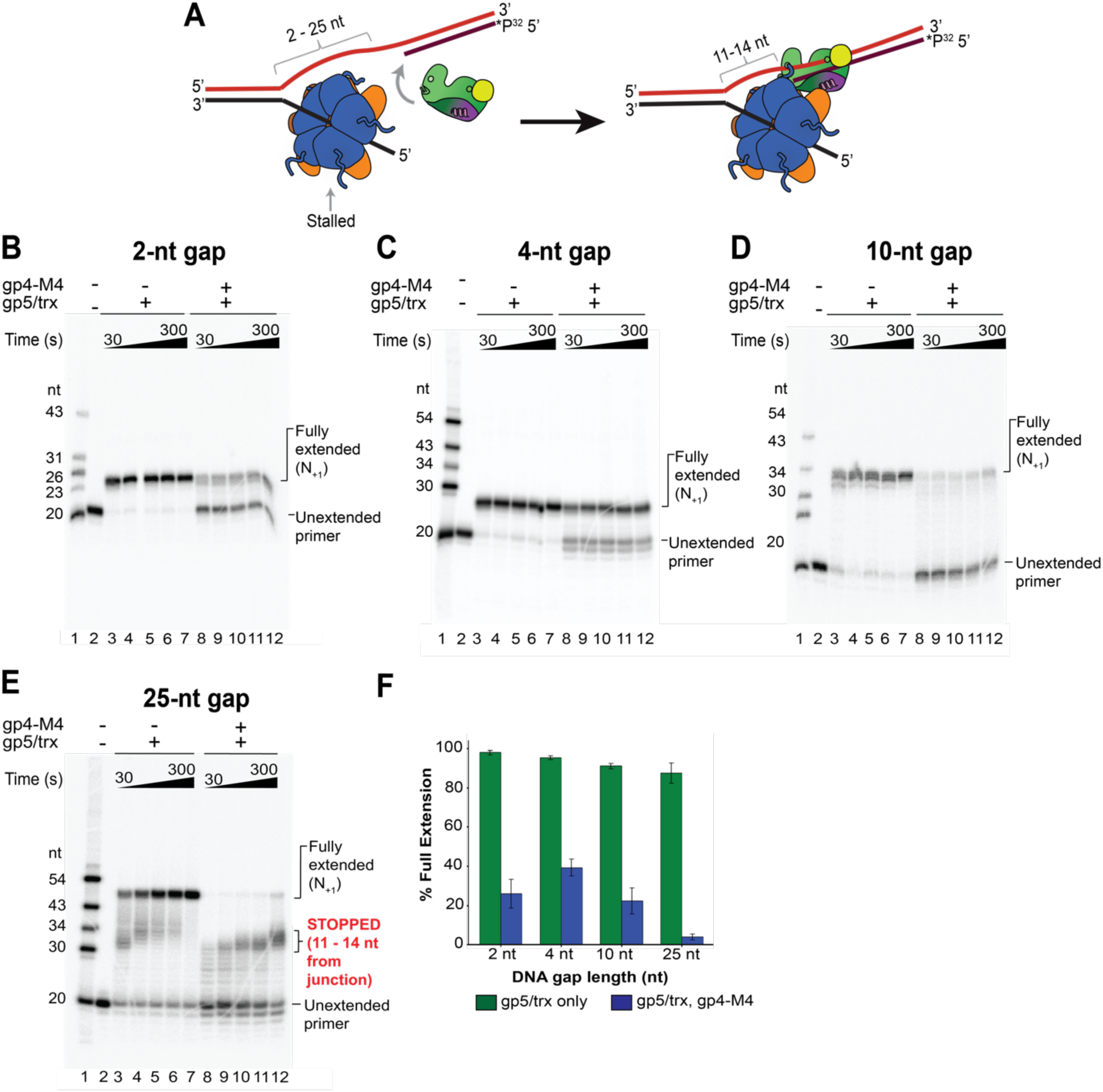
The stop-trap assay defines the ssDNA spacing on the leading-strand required for functional interactions between DNA primase-helicase and DNA polymerase. (**A**) Stop-trap experimental design. DNA fork substrates containing ssDNA gaps of 2-25 nt between the fork junction and the primer-template were assembled by annealing leading-strand oligonucleotides of varying length (35-58 nt) to a 40-nt lagging-strand and 20-nt primer (5′-end labeled with ³²P). Gp4-M4 was first preassembled on the DNA fork by incubating 0.83 µM gp4-M4 (hexamer concentration) with 0.3 µM DNA fork substrates in the presence of 0.1 mM dTTP at 37 °C for 10 minutes. DNA synthesis was initiated by the addition of 0.3 µM gp5/trx together with 0.1 mM each of dCTP, dGTP, and chain-terminating ddATP to halt strand displacement 2 bp inside the parental duplex. Reactions were stopped at indicated time points (30, 60, 90, 120, 300 s) with the addition of 20 mM EDTA. The schematic illustrates that only longer gaps (25 nt) support the formation of a functional complex in which gp4-M4 stalls gp5/trx approximately 11-14 nt from the fork junction. (**B-E**) The DNA products were resolved on 8M, 20% polyacrylamide gels. Faint bands migrating below the unextended primer (20 nt) likely represent a truncated primer species not removed during PAGE purification. (**F**) Quantification of full-length product formation at 300 s, calculated as the ratio of the intensity of the full-length product band to the total intensity of all substrate and product bands. Error bars represent standard deviation of two independent experiments.

These results indicate that gap lengths of <10 nt are too short to accommodate both gp5/trx and gp4-M4 simultaneously at the replication fork and therefore cannot efficiently support leading-strand synthesis via the PDU model. In contrast, when the gap size is 25 nt, gp5/trx is halted through interactions with gp4-M4, positioning its polymerase active site approximately 11-14 nt away from the fork junction, and supporting the HDU model of leading-strand synthesis.

### W579 Stabilizes DNA Polymerase on the DNA template

One of the key features of the PDU model is the proposed role of the P′ loop in gp5/trx, particularly W579 located within this loop, to act as an intercalating wedge that separates base-pairs and unwinds dsDNA (**Fig. 1B** and **SI Appendix**, **Fig. S2**), (4). Nevertheless, it has been previously shown that gp5/trx can only displace 4-5 bp before stalling as measured in rapid quench-flow assays (44) and up to 40-90 bp as reported in ensemble strand-displacement assays where multiple binding and strand-separation events can occur on one DNA ligand (45). Thus, the strand-displacement activity of gp5/trx is insufficient to unwind the approximately 16,000 bp genome of bacteriophage T7. In addition, combining dsDNA unwinding and primer synthesis activities on the leading-strand implies a 2-nt gap between the 3′ end of the primer and the replication fork junction (**Fig. 1B**). However, DNA primase-helicase binds weakly to DNA forks containing gaps of fewer than 10 nt (**Fig. 4 B** and **C**) and such DNA ligands fail to support the specific DNA polymerase-DNA primase-helicase interactions on the leading-strand (**Fig. 6B-D**). Therefore, to further clarify the role of the P′ loop, we constructed an exonuclease-deficient gp5/trx mutant in which W579 was substituted with alanine (gp5-W1/trx) and examined its DNA polymerization, strand-displacement, and DNA binding activities.

We first assessed the polymerization activity of gp5/trx and gp5-W1/trx. We measured the ability of gp5 variants to perform primer extension using a 58-nt template strand annealed to a 20-nt primer labeled at the 5’ end with ^32^P (**SI Appendix**, **Fig. S7A**). Reactions were carried out for up to 2 minutes with equimolar amounts of protein and DNA (0.3 µM), 10 mM MgCl₂, consistent with the concentration used previously (16, 19, 24, 46–54), and two different dNTP concentrations. Given the physiological concentration of all four dNTPs in *E. coli* is approximately 0.5 mM (55), we included a condition reflecting near-physiological dNTP levels (a total of 0.4 mM; 100 µM each nucleotide; ∼1333.3-fold molar excess over protein). Because recent studies suggested that polymerization activities of gp5/trx may be dependent on dNTP concentration (52), a high concentration of dNTPs was also used (a total of 7.2 mM; 1.8 mM each nucleotide; 24,000-fold molar excess over protein). Reactions were stopped by the addition of 20 mM EDTA, and the resultant DNA products were analyzed by denaturing PAGE (**SI Appendix**, **Fig. S7 B** and **C**). The sequencing gels revealed that at the lower dNTP concentration, gp5/trx efficiently extended primers to full-length products, corresponding to 58 nt (**SI Appendix**, **Fig. S7B**, lanes 7-10). In contrast, gp5-W1/trx exhibited reduced activity, with an approximately 3-fold decrease in yield and 12-fold decrease in extension rate (**SI Appendix**, **Fig. S7B**, lanes 17-20; quantified in **Fig. S7D**). Additionally, a significant fraction of products were intermediate in-length, ranging from 25 nt to 40 nt (**SI Appendix**, **Fig. S7B**, lanes 17-20). Interestingly, a substantial fraction of reaction products were not denatured upon addition of 8M urea. Nevertheless, these fragments represent fully extended products, as confirmed by including 58-nt ssDNA and 58-bp dsDNA controls on the gel (**SI Appendix**, **Fig. S7B**, lanes 1-3). Notably, at the high dNTP concentration, the polymerization activities of gp5-W1/trx and gp5/trx are similar (**SI Appendix**, **Fig. S7C**, lanes 3-6 and 13-16, respectively), consistent with previous findings (52).

In the following experiment, we assessed the strand-displacement activity of gp5/trx and gp5-W1/trx. A DNA fork substrate was assembled by annealing 43-nt and 40-nt ssDNA oligonucleotides corresponding to the leading- and lagging-strands, respectively, with a 20-nt 5′-³²P-labeled leading-strand primer (**SI Appendix**, **Fig. S7E**). Reactions were carried out under similar conditions described above. The sequencing gels revealed that gp5/trx fully extends the 20-nt primer and displaces the 13-bp region, resulting in 43-nt products on the gel (**SI Appendix**, **Fig. S7F**, lanes 3-6). Similar to the previous experiment, a substantial fraction of fully extended dsDNA reaction products were not denatured upon addition of 8M urea (**SI Appendix**, **Fig. S7F**). In contrast, gp5-W1/trx extends the primer approximately to the fork junction at the lower dNTP concentration but does not displace the lagging-strand (**SI Appendix**, **Fig. S7F**, lanes 13-16). Under the high dNTP concentration, gp5-W1/trx is able to carry out strand-displacement, as demonstrated by the presence of 43-nt ssDNA and 43-bp dsDNA fragments on the gel (**SI Appendix**, **Fig. S7G**, lanes 13-16), albeit at nearly 1.5-fold decreased yield and 5-fold reduced rate, as compared to gp5/trx (**SI Appendix**, **Fig. S7H**). Varied polymerization and strand-displacement activities of gp5-W1/trx at different dNTP concentrations suggest that W579 from the P′ loop may contribute to DNA binding, as the protein associates more readily with DNA at high non-physiological dNTP concentrations.

We have thus employed EMSA to assess the DNA-binding affinity of the gp5/trx variants (**Fig. 7** and **SI Appendix**, **Fig. S8**). The DNA ligand used in this experiment was analogous to that used in the polymerization assay (**SI Appendix**, **Fig. S7A**), except ddCTP was incorporated at the 3’-end of the 20-nt primer to prevent extension by DNA polymerase (**Fig. 7A**). DNA binding was assessed at both low (0.1 mM) and high (1.8 mM) concentrations of the next incoming nucleotide, dTTP, as described above. Gp5/trx binds to the primer-template with the K_d_ of 0.11 ± 0.02 µM (**Fig. 7B**). In contrast, gp5-W1/trx exhibits substantially reduced binding affinity, with approximately 10-fold weaker binding at the low dTTP concentration and 6-fold weaker binding at the high dTTP concentration (K_d_ > 0.9 µM and 0.59 ± 0.11 µM, respectively; **Fig. 7B**). Increased concentration of the next incoming nucleotide may help compensate for the reduced DNA binding affinity of gp5-W1/trx by promoting a closed conformation of the fingers domain in DNA polymerase, thereby stabilizing the polymerase on the DNA and increasing its polymerization and strand-displacement activities, although to substantially lower levels than those observed for gp5/trx (**SI Appendix**, **Fig. S7**). In addition, gp5-W1/trx displayed low binding affinity for DNA fork ligands with different ssDNA gap sizes (0, 2, and 10 nt) between the 3’-end of the primer and fork junction (K_d_ > 0.9 µM for all gap sizes; **SI Appendix**, **Fig. S9**).

**Fig. 7.**
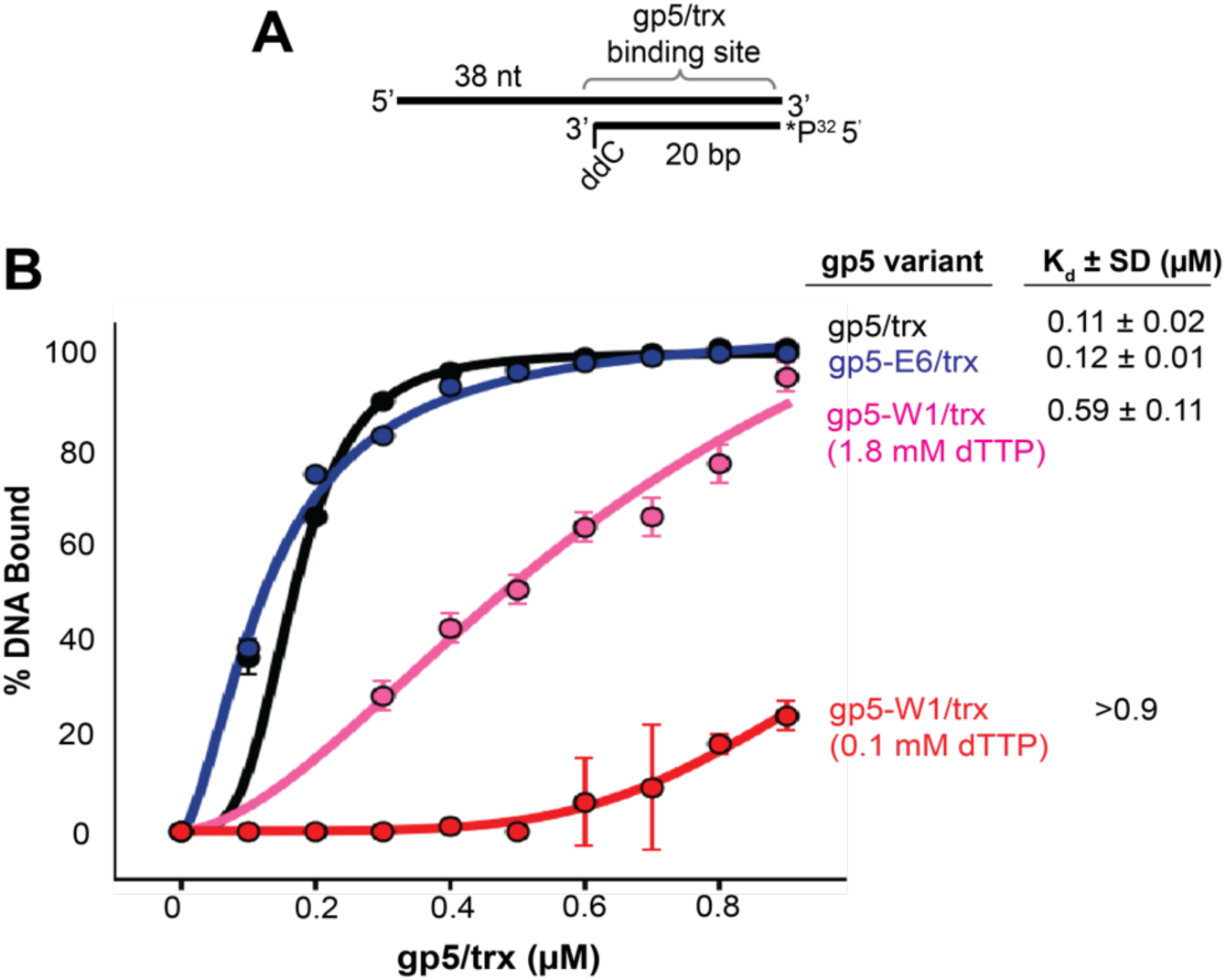
Characterization of DNA binding affinity of gp5/trx mutants. (A) DNA binding affinity of gp5/trx, gp5-E6/trx, and gp5-W1/trx was measured using a primer-template substrate obtained by annealing a 58-nt template strand with a 20-nt primer labeled with ³²P at the 5’ end. To prevent primer extension, the 3′ end of the primer contains a dideoxycytidine (ddC). The primer-template substrate (0.3 µM) was incubated with increasing amount of each gp5 variant (0.1-0.9 µM) and 0.1 mM of the next incoming nucleotide, dTTP, for 10 minutes at 37 °C. As we observe the strand-displacement and polymerization activities of gp5-W1/trx are largely dependent on dNTP concentration, we measured DNA binding affinity of gp5-W1/trx in the presence of both 0.1 mM and 1.8 mM dTTP. (**B**) Quantification of EMSA results of gp5/trx (black), gp5-E6/trx (blue), and gp5-W1/trx in the presence of 0.1 mM (red) and 1.8 mM (pink) dTTP. Dissociation constant (K_d_) is indicated next to each protein variant. Error represents standard deviation of three independent experiments.

In support of the HDU model, these results collectively indicate that W579 plays a role in DNA binding, rather than directly facilitating dsDNA unwinding, as proposed by the PDU model for the leading-strand replisome. A recent crystal structure of gp5/trx in complex with a primer-template and biochemical studies support this notion by showing that W579 forms a stacking interaction with the template base located 2 nt downstream from the polymerization active site (52), (**SI Appendix**, **Fig. S2D**). This interaction likely helps guide the template strand into the DNA polymerase active site after gp4 primase-helicase has separated the parental duplex DNA.

### Mutational Analysis of the Protein-Protein Interactions in the Leading-strand Replisome

The analysis of the HDU and PDU models identified the α-helix E* (residues Glu126-Gln150) from the exonuclease domain of gp5/trx as a structural feature that distinguishes the two models due to its mutually exclusive interactions with gp4 (**Fig. 2**). In the HDU, this α-helix is located at the unique interface formed by the leading-strand DNA polymerase and DNA primase-helicase (**Fig. 2A**), (3). In contrast, according to the PDU model, α-helix E* does not participate in interactions with DNA primase-helicase and remains solvent-exposed, positioned approximately 10 nm away from DNA primase-helicase (**Fig. 2B**), (4). The α-helix E* contains six acidic residues (Glu134, Glu138, Asp141, Asp142, Glu148, and Glu149) that we refer to as the acidic patch (Ap; **Fig. 2C**, inset). To investigate the interactions within the leading-strand replisome, we constructed an exonuclease-deficient gp5 variant (D5A and D7A) in which the six acidic residues of the Ap were replaced with alanines (gp5-E6).

Because α-helix E* is located within the exonuclease domain, we purified an exonuclease-positive DNA polymerase variant (gp5-E6exo^+^/trx) to investigate if the Ap mutations interfere with the proofreading activity of the exonuclease domain. The results of the exonuclease assay performed with a 54-nt ssDNA labeled with ^32^P at the 5’-end show that the exonuclease activity of gp5-E6exo^+^/trx is comparable to the activity of wild-type gp5/trx (gp5-exo^+^/trx), (**SI Appendix**, **Fig. S10**, lanes 8-11 and 3-6, respectively). In contrast, gp5-E6/trx does not display any exonuclease activity in the assay (**SI Appendix**, **Fig. S10**, lanes 13-16). Next, we examined the DNA polymerization (**SI Appendix**, **Fig. S7A-D**), strand-displacement (**SI Appendix**, **Fig. S7E-H**), and DNA binding (**Fig. 7**) activities of gp5-E6/trx using assays describe in the previous section concerned with gp5-W1/trx. Gp5-E6/trx and gp5/trx display a nearly identical polymerization and strand-displacement activities measured at two different dNTP concentrations (**SI Appendix**, **Fig. S7**) and DNA binding affinities to a primer-template (**Fig. 7**) with the dissociation constants of 0.12 ± 0.01 µM and 0.11 ± 0.02 µM, respectively (**Fig. 7B**).

Having established that the six amino acid substitutions in the α-helix E* do not affect the catalytic activities of DNA polymerase, we employed the thermal shift assay (TSA) to assess how they affect the ability of DNA polymerase to form a stable leading-strand replication complex (**Fig. 8**). The TSA detects changes in protein thermal stability by monitoring shifts in melting temperature (Tₘ), measured at the midpoint of the melting transition, that reflect stabilization upon binding to DNA or protein (56, 57). SYPRO Orange dye was used to monitor protein unfolding, as it fluoresces upon binding to hydrophobic residues exposed during thermal denaturation. In parallel with gp5-E6/trx, we also investigated gp5/trx and gp5-W1/trx.

**Fig. 8.**
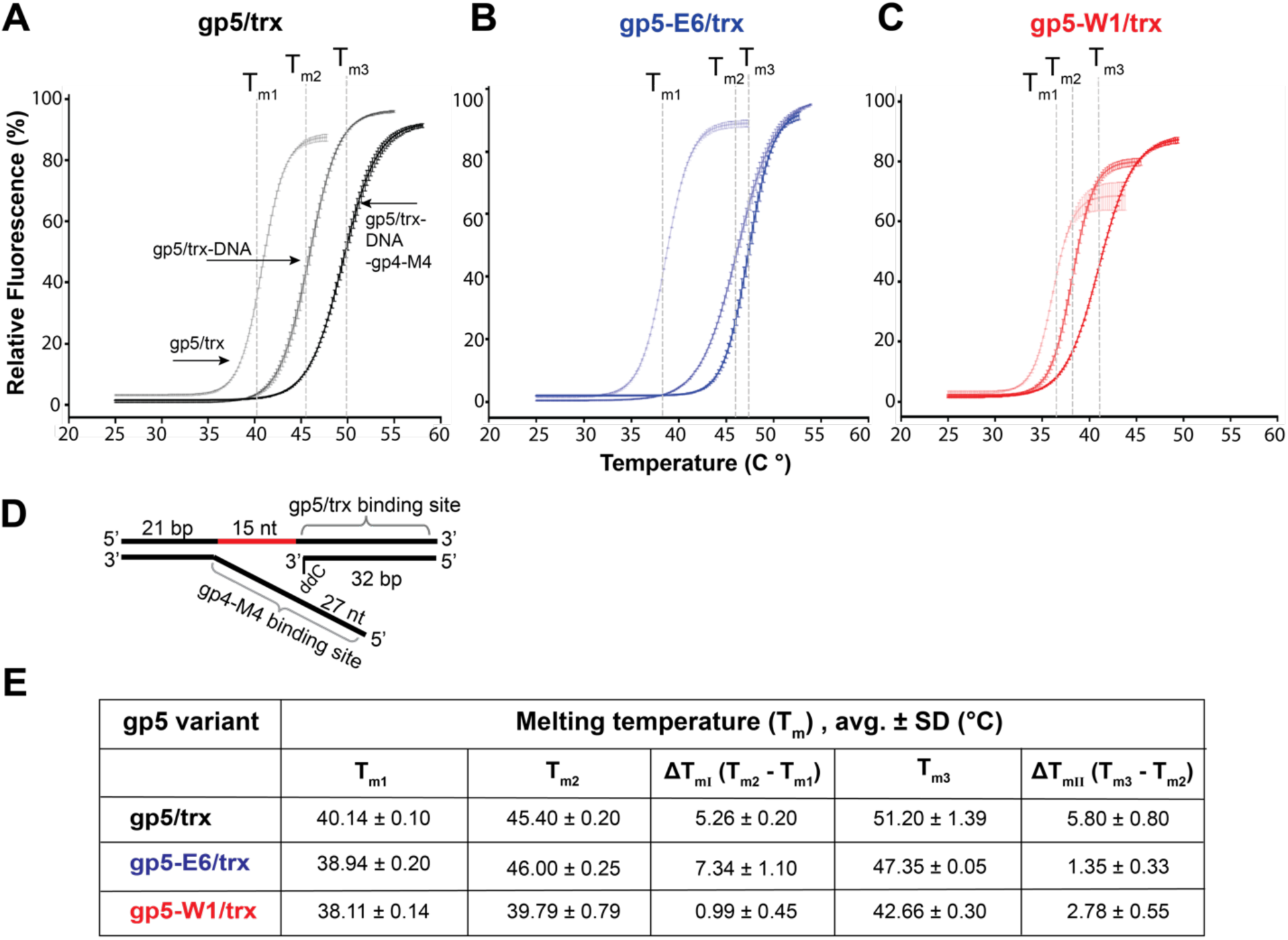
Thermal shift analysis shows that mutations in E* helix of gp5/trx disrupt formation of a stable leading-strand complex. Thermal stability of gp5/trx variants was monitored by SYPRO Orange fluorescence, which increases upon exposure of hydrophobic residues during protein denaturation. Reactions containing 1 µM gp5/trx variant (gp5/trx, gp5-E6/trx, or gp5-W1/trx) were heated gradually, and fluorescence was recorded as a function of temperature. Melting temperatures (Tₘ) were determined from the peak of the first derivative of the fluorescence signal with respect to temperature, corresponding to 50% unfolding. (**A-C**) Representative melting curves for gp5/trx (**A**), gp5-E6/trx (**B**), and gp5-W1/trx (**C**) under three conditions: protein alone (T_m1_), in the presence of a DNA fork with a 15-nt ssDNA gap (T_m2_), and with the DNA fork pre-bound by gp4-M4 (T_m3_). Error bars represent standard deviation of three independent experiments. (**D**) Schematic of the DNA fork substrate with 15-nt ssDNA gap. (E) Summary table of melting temperatures (T_m_) and changes in melting temperatures (ΔT_m_) for each condition. ΔT_mI_ represents the shift upon gp5/trx binding the DNA fork (T_m2_ - T_m1_), and ΔT_mII_ represents the additional shift associated with complex formation upon addition of gp5/trx to the pre-bound gp4-M4-DNA complex (T_m3_ - T_m2_).

We first measured the Tₘ of gp5/trx variants (Tₘ_1_). Samples containing 1 µM of each gp5/trx variant and SYPRO Orange dye were gradually heated and changes in fluorescence were monitored. Gp5-E6/trx and gp5-W1/trx exhibited similar Tₘ_1_ to gp5/trx (38.94 ± 0.2 °C, 38.11 ± 0.14 °C and 40.14 ± 0.1 °C, respectively), suggesting that the solvent-exposed mutations did not disrupt the overall protein fold in solution (**Fig. 8A-C**). Next, we evaluated Tₘ of the gp5 variants in the presence of a DNA fork ligand (Tₘ_2_) containing a 15-nt ssDNA gap between the 3’-end of the primer and fork junction (**Fig. 8D**). A 15-nt gap was chosen to reflect the condition required to form a stable leading-strand replication complex, as established in the previous section (**Fig. 6E**).

We measured changes in melting temperatures (ΔTₘ_I_ = Tₘ_2_ - Tₘ_1_) at several DNA:protein molar ratios, ranging up to 5-fold excess DNA (see **Materials and Methods**). The data presented in **Fig. 8** reflect the optimal 1:1 ratio, which produced the maximal shift, as further addition of DNA to reaction mixtures did not affect the measured Tₘ. The ΔTₘ_I_ observed for gp5/trx (5.26 ± 0.20 °C) and gp5-E6/trx (7.34 ± 1.1 °C) indicate that DNA polymerase is stabilized upon binding to the DNA fork, (**Fig. 8A** and **B**, respectively). In contrast, gp5-W1/trx exhibited only a minimal ΔTₘ_I_ of 0.99 ± 0.45 °C (**Fig. 8C**). Shifts in Tₘ of less than 2 °C are generally considered not significant (56–58), as they fall within the typical experimental variability of the TSA and could arise from minor differences in protein concentration, dye binding, or heating rate. Thus, the shift observed for gp5-W1/trx in the presence of DNA does not reflect meaningful stabilization. This result is consistent with the poor DNA binding affinity of gp5-W1/trx revealed by EMSA in the previous section (**Fig. 7B** and **SI Appendix**, **Fig. S8**, **S9**).

To assess how the protein-protein interactions influence the stability of the leading-strand replisome, we measured the Tₘ of gp5/trx variants in the presence of a DNA fork with pre-bound gp4-M4 (Tₘ_3_). The optimal stoichiometry of gp4-M4:DNA:gp5/trx variants (6:2:1) was established by employing the protocol described above. We first mixed gp4-M4 and the DNA fork at different molar ratios to achieve the maximal shift in melting temperature, beyond which further addition of DNA does not affect the result. These experiments revealed the 6:2 stoichiometry of gp4-M4:DNA binding and indicate that nearly all gp4-M4 molecules are involved in the formation of hexameric protein complexes with DNA, reflecting the physiological relevance of the TSA measurements. Gp5/trx variants were then titrated to the reaction mixtures in a similar fashion (see **Materials and Methods**). The results show that gp5/trx formed a specific complex with gp4-M4 and the DNA fork, as indicated by its increased stability reflected by a ΔTₘ_II_ (ΔTₘ_II_ = Tₘ_3_ - Tₘ_2_) of 5.8 ± 0.8 °C (**Fig. 8A**). In contrast, gp5-E6/trx showed only a minimal change in melting temperature (ΔTₘ_II_ = 1.35 ± 0.33 °C), which falls within the experimental error of the method, demonstrating that although gp5-E6/trx binds to the DNA fork, the amino-acid mutations within the Ap in α-helix E* prevent its binding to the DNA primase-helicase (**Fig. 8B**). Likewise, the addition of gp5-W1/trx to gp4-M4-DNA samples did not significantly change ΔTₘ_II_ (2.87 ± 0.55 °C), demonstrating that the impaired DNA-binding affinity of gp5-W1/trx prevents the formation of a stable leading-strand replication complex (**Fig. 8C**). A summary of all measured Tₘ and ΔTₘ values is presented in **Fig. 8E**.

Taken together, the results of the mutational analysis strongly support the HDU model, as altering the DNA polymerase-DNA primase-helicase interface identified in the cryo-EM structure by Kulczyk et al. (3) prevents the formation of the leading-strand replisome. In contrast, the results argue against the PDU model (4), in which the solvent-exposed α-helix E* in DNA polymerase makes no interactions with DNA primase-helicase. If this model were correct, the mutations introduced in α-helix E* would have no effect on the assembly of the leading-strand replisome.

### Processive Leading-Strand DNA Synthesis Requires Interactions Between DNA Polymerase α Helix E* and the Primase Domain of DNA Primase-Helicase

Having established the importance of the interaction between the α-helix E* from DNA polymerase with DNA primase-helicase for leading-strand replisome assembly, we examined whether gp5-E6/trx and other DNA polymerase variants (gp5/trx, gp5-W1/trx) can support processive leading-strand synthesis using a DNA minicircle assay (59), (**Fig. 9**). Reaction mixtures contained 80 nM gp5/trx variant, 100 nM minicircle, 10 nM WTgp4 (hexameric concentration), and a ^32^P-labeled deoxyguanosine triphosphate ([α-³²P]-dGTP). The minicircle substrate consisted of a 70-nt circular ssDNA annealed to a 110-nt ssDNA, forming a structure that mimics a replication fork, with a 40-nt 5′-overhang that serves as a loading site for gp4 (**Fig. 9A**), (59). Gp5/trx performed processive leading-strand synthesis, as evidenced by efficient incorporation of [α-³²P]dGMP into nascent DNA, reaching a plateau after 30-minute incubation at 707.44 ± 56.06 picomoles corresponding to an average processivity of 15.4 kb ± 0.18 kb (**Fig. 9B**). Alkaline agarose gel electrophoresis confirmed this result by revealing DNA product lengths ranging from approximately 3 kb to 20 kb (**SI Appendix**, **Fig. S11A-E**, lanes 2-4).

**Fig. 9.**
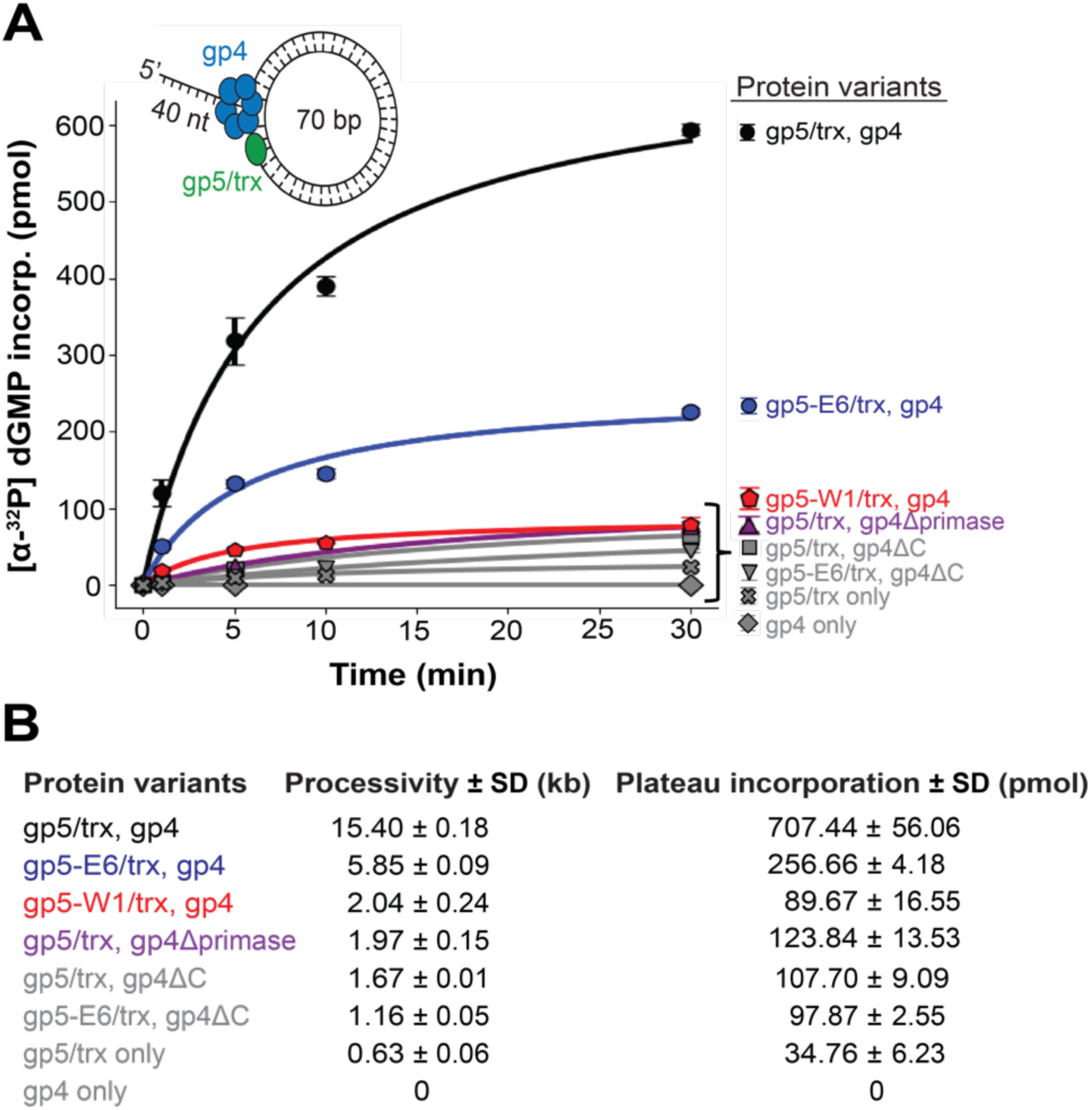
Characterization of leading-strand synthesis of gp5/trx and gp4 variants by minicircle assay. (**A**) Reactions contained 100 nM minicircle, 10 nM gp4 variant (hexameric concentration), 80 nM gp5/trx variant, 1.8 mM of each dNTP, and [α-^32^P]dGTP at 600 mCi/mmol to measure synthesis. Reactions were incubated at 37°C for the indicated times (1, 5, 10, 30 min) and quenched by addition of 20 mM EDTA. DNA synthesis was quantified by spotting reactions onto DE81 filter paper and measuring [α-^32^P]dGMP incorporation by liquid scintillation counter as described in the **Materials and Methods**. Error bars represent standard deviation of three independent trials. (**B**) Table of maximum dGMP incorporation (plateau) and processivity, obtained from **A**, for protein variants.

Importantly, in a control experiment with gp5/trx in the absence of WTgp4, minimal nucleotide incorporation into the minicircle was detected (34.76 ± 6.23 picomoles corresponding to an average processivity of 0.63 kb ± 0.06 kb), (**Fig. 9A** and **B**). This result indicates that DNA polymerase alone cannot support leading-strand synthesis and argues strongly against the PDU model. Likewise, no signal was observed in a control reaction containing WTgp4 only, where [α-³²P]-dGTP could not be incorporated into DNA (**Fig. 9B**).

In contrast, gp5-E6/trx exhibited an approximately 65% reduction in DNA synthesis, reaching a plateau at 256.66 ± 4.18 picomoles and an average processivity of 5.85 kb ± 0.09 kb (**Fig. 9B**). Given that gp5/trx and gp5-E6/trx display similar DNA polymerization and strand-displacement activities (**SI Appendix**, **Fig. S7**), comparable DNA-binding affinities (**Fig. 7**), and produce analogous DNA products in the minicircle reaction ranging from ∼3 kb to 20 kb (**SI Appendix**, **Fig. S11A**, lanes 2-4 and 5-7, respectively), these results indicate that the reduction in leading-strand synthesis by gp5-E6/trx arises from the loss of an essential interaction between the E* helix of gp5 and gp4 that stabilizes the processive leading-strand replisome.

In the HDU model of the leading-strand replisome, the α-helix E* makes contacts with the primase domain of gp4. To investigate the reciprocal interaction on gp4, we generated a truncated variant of gp4 lacking the primase domain (gp4Δprimase) and measured its ability to support leading-strand synthesis in the minicircle reaction (**Fig. 9 A** and **B**). Gp4Δprimase displayed an approximately 90% reduction in DNA synthesis, reaching a plateau at 123.84 ± 13.53 picomoles and an average processivity of 1.97 ± 0.15 kb, confirming the role of the primase domain in processive leading strand synthesis, and consistent with previous single-molecule experiments (24). DNA products ranged from 3 kb to 20 kb, as revealed by alkaline agarose gel electrophoresis (**SI Appendix**, **Fig. S11B**, lanes 5-7). In addition, to determine whether gp4Δprimase can form specific interactions with DNA polymerase during leading-strand synthesis, we constructed a gp4Δprimase variant containing an additional mutation (E343Q) in the helicase domain (gp4-M1Δprimase), which abolishes the protein’s ability to hydrolyze nucleotides and translocate along ssDNA. We investigated gp4-M1Δprimase in the stop-trap experiment with gp5/trx using a DNA fork substrate containing a 25-nt gap between the 3’-end of the primer and the fork junction (**SI Appendix**, **Fig. S12A**). Unlike gp4-M4 (**Fig. 6E**), gp4-M1Δprimase was unable to halt DNA polymerase, and all DNA products were fully extended position N_+1_ within the first 30 s of the reaction (**SI Appendix**, **Fig. S12B**, lanes 14-18), further confirming the role of the primase domain in its reciprocal interaction with gp5/trx.

Gp5-W1/trx revealed a 90% reduction in total dNTP incorporation over 30 minutes with a plateau at 89.67 ± 16.55 picomoles and an average processivity of 2.04 ± 0.24 kb, as compared to gp5/trx (**Fig. 9B**). In addition, leading-strand DNA synthesis produced shorter products ranging from ∼2 kb to 5 kb (**SI Appendix**, **Fig. S11C**, lanes 5-7). These findings are consistent with the decreased DNA polymerization and strand-displacement activities (**SI Appendix**, **Fig. S7**), and the reduced DNA-binding affinity of gp5-W1/trx (**Fig. 7** and **SI Appendix**, **Fig. S8**, **S9**). We hypothesize that the weak DNA binding likely causes gp5-W1/trx to dissociate from the template prematurely, resulting in truncated products. Importantly, gp5-W1/trx supports only limited leading-strand synthesis (**Fig. 9 A** and **B**), highlighting the role of W579 and the P′ loop in DNA binding rather than in the unwinding of dsDNA. The results of the minicircle (**Fig. 9** and **SI Appendix**, **Fig. S11**) and stop-trap experiments (**Fig. 6**), together with the TSA measurements of the leading-strand replisome assembly presented in the previous section (**Fig. 8**), further support the HDU model for leading-strand replication.

### Deletion of the DNA Primase-Helicase C-Terminal Tails Reduces Processivity but Does Not Abolish Processive Leading-Strand Synthesis

According to the PDU model, the interaction between DNA polymerase and DNA primase-helicase is solely mediated through the C-terminal tails of gp4 (4). In contrast, although the interaction between the acidic C-terminal tail of gp4 and the Fbp of gp5/trx may be maintained in HDU, it is not essential for the stability of the leading-strand replisome (**Fig. 1A**). Therefore, to investigate the role of this interaction in processive leading-strand synthesis, we generated a gp4 mutant lacking the 17 residue C-terminal tail (gp4ΔC). First, in a control experiment, we tested the DNA unwinding activity of gp4ΔC using the DNA substrate obtained upon annealing a 70-nt minicircle and a 110-nt ssDNA oligonucleotide with a 5’ ^32^P label (**SI Appendix**, **Fig. S13A**). In this assay, the ^32^P labeled-strand is released as ssDNA upon helicase-mediated unwinding, and the resulting ssDNA and dsDNA species are separated by nondenaturing PAGE (**SI Appendix**, **Fig. S13B-D**). Gp4ΔC unwound dsDNA at approximately 45% of the rate observed for WTgp4 (**SI Appendix**, **Fig. S13E**), indicating a role for the negatively charged C-terminal tails of gp4 in DNA unwinding and consistent with previous studies (24). Next, we tested gp4ΔC with gp5/trx in a minicircle reaction (**Fig. 9** and **SI Appendix**, **Fig. S11D**). Although DNA synthesis was reduced to ∼15% of WTgp4 activity (plateau at 107.7 ± 9.09 picomoles and an average processivity of 1.67 ± 0.01 kb), gp4ΔC supported leading-strand synthesis with DNA products ranging from 1.5 kb to 5 kb as revealed by alkaline gel electrophoresis (**SI Appendix**, **Fig. S11D**, lanes 5-7), inconsistent with the PDU model. In an analogous reaction performed with gp4ΔC and gp5-E6/trx, no DNA products were detected on the alkaline gel (**SI Appendix**, **Fig. S11E**, lanes 5-7), highlighting the importance of the interaction between the α-helix E* from DNA polymerase with DNA primase-helicase.

Our findings are consistent with previous studies (20, 24–27), confirming that gp4ΔC efficiently assembles into the leading-strand replisome and supports leading-strand synthesis. Surface plasmon resonance binding experiments with gp5/trx bound to a primer-template showed that deletion of the C-terminal tail of gp4 reduces gp4ΔC binding to 68 ± 9% of the level observed for the WTgp4, but does not lead to dissociation of the leading-strand complex (25).

Notably, flow-stretching single-molecule experiments using DNA forks confirmed that gp4ΔC can support leading-strand replication, albeit with a lower rate and processivity compared to WTgp4 (90 ± 9 bp/s and 4 ± 0.8 kb for gp4ΔC versus 112 ± 24 bp/s and 16 ± 4 kb for WTgp4), (24). Approximately a 75% reduction in processivity is particularly significant, as it indicates that the interaction between DNA polymerase and the C-terminal tail of gp4 contributes to processivity by tethering polymerases that transiently dissociate from the primer-template during ongoing leading-strand replication. This occurs during polymerase exchange, a process we previously directly visualized by correlating the stoichiometry of individual fluorescently labeled T7 DNA polymerases at the replication fork with DNA synthesis using a single-molecule assay that combines DNA flow-stretching with total internal reflection fluorescence microscopy (60). This interaction is mediated by the TBDbp region of gp5/trx (25, 27), as elimination of the basic charges in the exchange patch also reduces the processivity of leading-strand synthesis to 6 ± 3 kb (25). We observed that at saturating concentrations of gp5/trx, all C-terminal binding sites on gp4 are occupied by DNA polymerases during ongoing DNA synthesis, including the enzyme associated with the primer-template and five additional DNA polymerases associated with the fork. The excess DNA polymerases remain passively associated with the replisome through electrostatic interactions involving TBDbp and the C-terminal tails of DNA primase-helicase for approximately 50 seconds until a stochastic and transient dissociation of the synthesizing enzyme from the primer-template allows for a polymerase exchange to occur (27). These results collectively underscore the necessity of additional interactions between gp5/trx and gp4 implied by the HDU model and suggest that the DNA polymerase-DNA primase-helicase arrangement depicted in the PDU model may represent the replisome conformation during transient interactions that occur during DNA polymerase exchange or loading (**Fig. 10**).

**Fig. 10.**
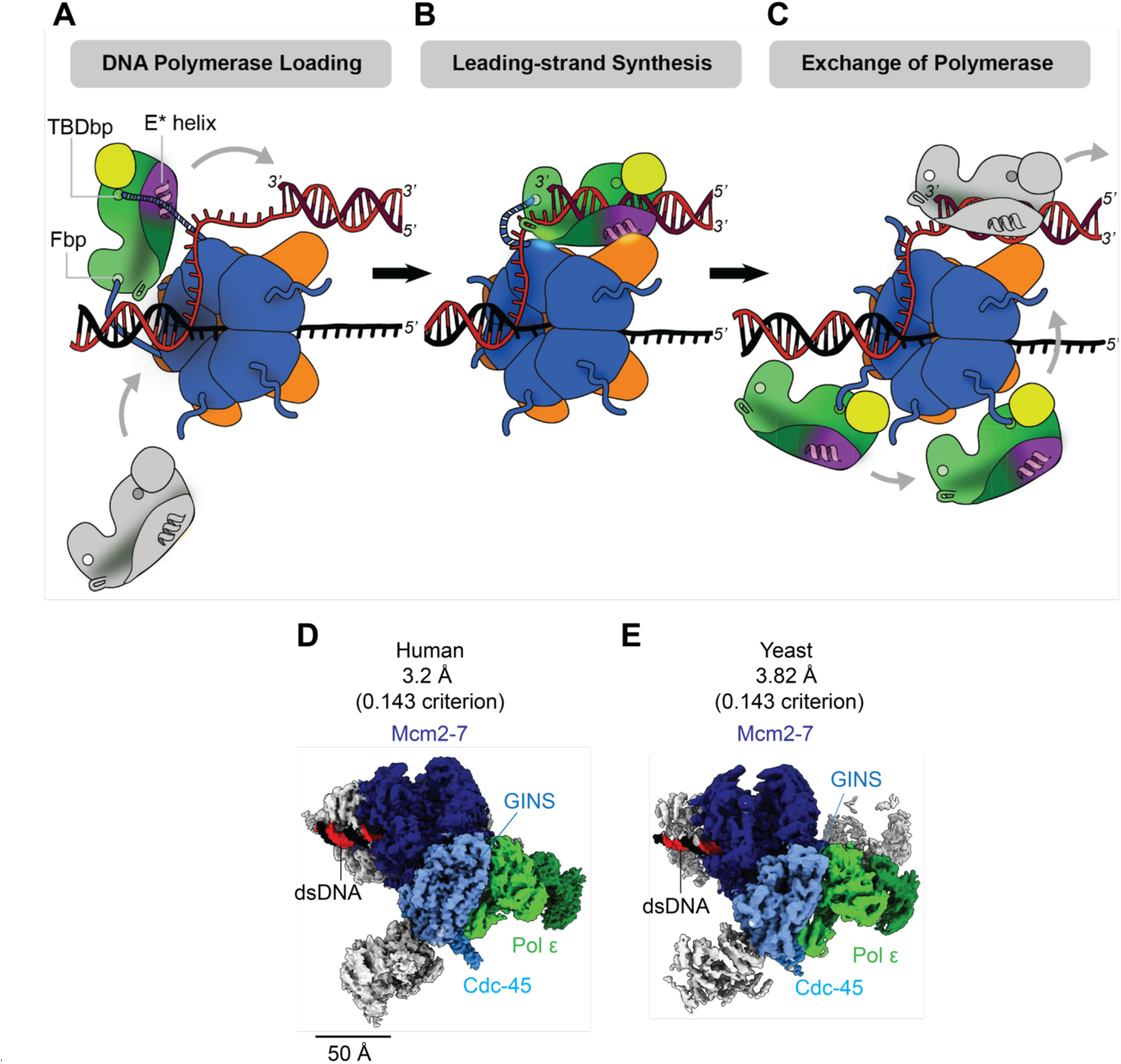
Leading-Strand Replication in Bacteriophage T7 and across taxa. (**A-C**) Catalytic cycle of T7 leading-strand synthesis. (**A**) gp4 assembles as a hexamer on the lagging-strand at the replication fork and recruits gp5/trx through interactions between the acidic C-terminal tails of gp4 and the front basic patch (Fbp) of gp5/trx. Gp4 may also engage the basic patch in the thioredoxin binding domain (TBDbp) of gp5/trx during recruitment; however, productive loading of gp5/trx onto the leading-strand primer-template requires binding to be switched to the Fbp. (**B**) During processive leading-strand synthesis, gp4 unwinds dsDNA at the fork junction, generating an 11-to 14-nt ssDNA template that is extended by the trailing leading-gp5/trx. The unique E* helix of gp5/trx forms stable interactions with the primase domain of gp4. Additional stabilizing contacts are formed between the adjacent helicase domain of gp4 and polymerase domain of gp5. Interactions between the C-terminal tails and the Fbp may also be maintained throughout synthesis to increase the processivity of the complex. (**C**) If gp5/trx dissociates from the primer-template, gp4 can facilitate re-loading through interactions involving the TBDbp to allow synthesis to resume. In addition, the multiple C-terminal tails of the gp4 hexamer can recruit additional gp5/trx molecules via either basic patch, potentially serving as a reservoir of polymerases at the replication fork. (**D-E**) Cryo-EM structures of leading-strand complexes from Homo sapiens (EMDB ID: 13375) (**D**) and Saccharomyces cerevisiae (EMDB: 52459) (**E**) support an HDU model of leading-strand replication in which the CMG (Cdc45-MCM2-7-GINS) helicase is positioned at the fork junction to unwind dsDNA for extension by leading-strand polymerase Pol ε. In the structures shown, MCM2-7 is colored navy, GINS light blue, Cdc45 cyan, Pol ε green, dsDNA black/red, and accessory proteins gray.

## Discussion

### Leading-Strand Replication in Bacteriophage T7

Bacteriophage T7 is a model system to study DNA replication. The catalytic cycle of DNA replication involves multiple steps and requires coordinated interactions between DNA primase-helicase and DNA polymerase. In phage T7, gp4 first loads onto lagging-strand as a hexamer. The leading-strand gp5/trx is recruited to the replisome and loaded to the primer-template through an interaction between the Fbp of gp5 and the acidic C-terminal tail of gp4 (20), (**Fig. 10A**). DNA polymerase loading in bacteria and eukaryotes, however, relies on clamp proteins that tether the DNA polymerase to the DNA template (2). Gp5/trx may also bind the C-terminal tails of gp4 through the TBDbp of gp5/trx, but must switch its binding to the Fbp to initiate DNA synthesis (20), (**Fig. 10A**). Following loading, gp4 and gp5/trx engage in a highly stable interaction that increases the processivity of leading-strand synthesis from 0.8 kb to 5 kb per binding event (20, 24–27), (**Fig. 10B**). Additionally, electrostatic interactions between the C-terminal tail of gp4 and the TBDbp of gp5/trx increase processivity from approximately 4-6 kb to 16 kb by capturing gp5/trx that transiently dissociate from DNA (25, 27), (**Fig. 10C**). The specificity of gp4-gp5/trx interactions is illustrated by the inability of gp4 to perform leading-strand synthesis in the presence of DNA polymerases from other systems, including bacteriophage T4 and *E. coli* (61). Additional interactions between gp4 and gp5/trx are formed during lagging-strand synthesis. Furthermore, single-stranded DNA-binding protein, gp2.5, plays an important role in synchronizing the events on leading- and lagging-strands through interactions with both gp4 and gp5/trx through its acidic C-terminal tails that can, for example, compete with gp4 for binding to gp5/trx (1, 62, 63).

### Structural and Biochemical Evidence Supporting the HDU Model

Due to the complexity and dynamic nature of DNA replication, the physical positioning of proteins at the replication fork have remained unclear. Two mutually exclusive models (**Fig. 1**) have been proposed for leading-strand synthesis based on structural and biochemical analyses that we define as: (1) helicase-driven unwinding (HDU), in which the helicase migrates ahead of the DNA polymerase and separates the parental duplex into leading- and lagging-strands for DNA synthesis, and (2) polymerase-driven unwinding (PDU), in which the leading-strand DNA polymerase itself drives DNA unwinding while simultaneously synthesizing the leading strand. In this model, the DNA helicase plays an auxiliary role in stabilizing the lagging-strand for replication by the lagging-strand DNA polymerase.

Despite multiple structures of T7 replisome subassemblies (3–5, 24), none have revealed the full path of the DNA fork (**SI Appendix**, **Fig. S1** and **Movie S1**). In cryo-EM reconstructions reported in 2019 depicting the PDU model, the absence of continuous DNA density may reflect the use of DNA substrates with a 2-nt ssDNA gap between the fork junction and primer-template, and the inability of the resultant DNA fork to support the formation of a functional leading-strand replisome (4). Our biochemical data indicate that short ssDNA gaps prevent gp4 loading (**Fig. 3A-C**) and cannot support the formation of a leading-strand complex (**Fig. 6**). Furthermore, on this DNA substrate, the only protein-protein interactions observed in the cryo-EM maps are contacts between the acidic C-terminal tails of gp4 and the Fbp and/or the TBDbp of gp5/trx, which mediate gp5/trx loading and polymerase exchange, respectively (**Fig. 10**).

Consistent with previous studies (24, 38), we observe that leading-strand synthesis up to 5 kb is maintained in gp4 mutants lacking the C-terminal tails (**Fig. 9** and **SI Appendix**, **Fig. S11D**), indicating that additional contacts between gp4 and gp5/trx must stabilize the leading-strand replisome. Taken together, these observations suggest that the cryo-EM structures reported by Gao et al. may have captured loading or exchange intermediates rather than the architecture of the processive leading-strand replisome (**Fig. 10A**). Notably, none of the structures deposited in the RCSB PDB by Gao et al. contains DNA primase-helicase bound to more than one copy of DNA polymerase. Therefore, by definition, these structures do not represent the fully assembled replisome, which must include at least two DNA polymerases, responsible for synthesizing the leading- and lagging-strands, and, as a result, further suggesting that the proposed model may not accurately reflect the physiological complex. Furthermore, the lack of continuous density representing a DNA fork raises the possibility that the reconstructions reflect proteins bound to two separate DNA forks, i.e., DNA polymerase associating with a primer-template fragment of one DNA ligand, and DNA primase-helicase binding to the ssDNA portion of another DNA ligand. This could explain the unusual architecture of the DNA in the PDU model (see **Mechanistic and Kinetic Constraints Favor HDU** below and **Fig. 1**) and is consistent with the results of our DNA-binding experiments (**Fig. 3**-**6**). Gao et al. proposed that W579 of the P’ loop of gp5 functions as a separation pin in a PDU model (4). This interpretation is based on the positioning of W579 on a 2-nt gap DNA substrate, where it stacks with the template base at the fork junction. However, none of the reported leading-strand cryo-EM maps contain density to which W579 or the P’ loop can be unambiguously assigned (**SI Appendix**, **Fig. S2 A** and **B**), raising questions whether this loop directly participates in DNA unwinding. A recently reported crystal structure of a gp5/trx-DNA complex shows W579 contacts the template strand 2 nt from the polymerization active site, suggesting the P’ loop is important in stabilization of gp5/trx on DNA (52), (**SI Appendix**, **Fig. S2D**). Notably, this crystal structure contains only a primer-template substrate and lacks a displaced lagging strand. Consistent with our biochemical data (**Fig. 7**, **Fig. 8**), these observations support the interpretation that the W579 stabilizes gp5/trx DNA binding, rather than functioning as a separation pin for duplex unwinding.

By contrast, the cryo-EM structure of the bacteriophage T7 replisome reported by Kulczyk et al. in 2017, in which the DNA primase-helicase associates with two DNA polymerases on a DNA substrate containing a 10-nt ssDNA gap, supports an HDU architecture (3). Despite lack of continuous density representing a DNA fork in the structure (**SI Appendix**, **Fig. S1**), we find that this spacing more closely reflects the architecture required for formation of a leading-strand complex (**Fig. 6**). Nonetheless, our biochemical experiments suggest that slightly larger gaps of ssDNA approximately 11-15 nucleotides are optimal for complex formation (**Fig. 4**, **Fig. 6**). In vivo, this exposed ssDNA may be protected by gp2.5, which binds ssDNA as a dimer with consistent footprint of approximately 12-14 nucleotides (64–66). Guided by this structure (3), we used targeted mutagenesis to identify a specific interaction between the gp5 exonuclease domain of DNA polymerase and the primase domain of DNA primase-helicase that is required for robust leading-strand synthesis (**Fig. 8**, **Fig. 9**). In support of an HDU model, we show that contacts between the E* helix of gp5 and primase domain of gp4 stabilize the leading-strand complex (**Fig. 8**, **Fig. 9**).

However, as the loss of these interactions do not completely abolish leading-strand synthesis (**Fig. 9** and **SI Appendix**, **Fig. S11A**), gp4 and gp5/trx likely maintain additional interactions to coordinate processive leading-strand synthesis. The cryo-EM structure determined by Kulczyk et al. reveals that the α-helix P of gp5 contacts the α-helix E (Thr402-Leu416) in the helicase domain of gp4 (**SI Appendix**, **Fig. 2C**), (3). This notion is also supported by a low-resolution small angle x-ray scattering (SAXS) structure of the leading-strand complex (24). Because residues at the tip of the E* helix (Lys144-Gly157) are important for DNA binding during exonucleolytic degradation (67), interactions between the E* helix and gp4 may play a role in regulating proof-reading during leading-strand synthesis.

### Mechanistic and Kinetic Constraints Favor HDU Model

Another important consideration in distinguishing between the HDU and PDU models is the relative rates and mechanical capabilities of gp4 and gp5/trx. Individually, these enzymes operate at markedly different speeds. Gp5/trx catalyzes primer extension at an average rate of approximately 220 nt/s and can perform limited strand-displacement synthesis, incorporating 4-5 nt before stalling in rapid quench-flow assays (44) and up to 90 nt in ensemble assays where multiple binding and strand-displacements events take place on one DNA substrate (45). In contrast, gp4 translocates along ssDNA at 130 nt/s (44) but unwinds duplex DNA more slowly at 15-30 bp/s (19). However, the leading-strand replisome moves much faster than either enzyme’s isolated dsDNA activity, achieving synthesis rates of up to 160 nt/s (26, 59).

Under a PDU scenario, gp5/trx would simultaneously drive duplex unwinding and synthesis at 160 nt/s, while gp4 translocates along the lagging ssDNA produced by the polymerase. In this configuration, the forward force generated by helicase nucleotide hydrolysis would need to be transmitted into forward motion of DNA polymerase, despite the two enzymes moving along DNA substrates oriented nearly perpendicular to one another and being connected only through one or two flexible C-terminal linkers (**Fig. 1B** and **SI Appendix**, **Fig. S1**). How such force transduction could be achieved in the absence of rigid structural coupling is unclear. Moreover, gp5/trx alone exhibits poor strand-displacement activity, and it has been shown gp4 can displace a stalled gp5/trx from DNA (68).

In contrast, an HDU model posits that gp4 actively unwinds dsDNA at the replication fork, with gp5/trx rapidly copying the newly exposed leading-strand. In this case, the overall rate of synthesis is limited by the unwinding speed of gp4, rather than the polymerization activity of gp5/trx alone. Single-molecule experiments support this interpretation, demonstrating that the unwinding rate of gp4 increases substantially in the presence of a non-replicative polymerase, reaching up to approximately 200 bp/s (68). Furthermore, we have shown that gp5/trx cannot overtake a stalled gp4 (**Fig. 6**), consistent with a model in which helicase leads replisome progression.

### Structural Support of HDU model Across Taxa

To our current understanding, helicases facilitate dsDNA unwinding in all characterized systems (69–76), (**Fig. 10 D** and **E**). In eukaryotes, high-resolution structures of the yeast and human leading-strand replisomes have been determined with partially visualized fragments of DNA forks, both of which support an HDU model (70–72). In the eukaryotic replisome, both the leading-strand polymerase (Pol ɛ) and the Cdc45-Mcm2-7-GINS (CMG) helicase complex binds to the leading-strand (**Fig. 10 D** and **E**). Although the first reported eukaryotic replisome structure proposed a PDU model (77), this interpretation was later shown to result from an incorrect assignment of CMG polarity on DNA (75). Subsequent structures revealed the opposite architecture, with CMG positioned at the fork junction and Pol ε trailing behind (72, 75). In the human leading-strand replisome, the MCM7 subunit of CMG directly contacts the fork junction to act as a separation pin, while Pol ε binds the C-terminal tier of the MCM ring and captures the newly unwound leading-strand after it exits the CMG central channel (72). Extensive contacts between the C-terminal lobe of Pol ε and the backside of CMG further support tightly coupled helicase-polymerase progression during leading-strand synthesis (72–74), (**Fig. 10 D** and **E**).

Despite structural differences among DNA polymerases across taxa, including elements unique to T7 such as the E* helix, the overall organization and functional coupling of helicase and polymerase are consistent with the HDU model. Structural information for bacterial and viral replisomes remains more limited, as only structures of individual proteins and protein-DNA complexes have been determined (78–80). Nonetheless, a recent structure of a viral SF3 helicase bound to DNA reveals that the helicase directly interacts with the fork junction (76). The T7 replisome has long been regarded as a model system for elucidating the mechanisms of DNA replication. Our findings support the universality of the HDU model across diverse replication systems.

## Materials and Methods

### Cloning, Expression and Purification Recombinant Proteins

The gp5/trx complex was purified as a 1:1 complex to homogeneity in either the wild-type form (gp5-exo⁺) (81) or with alanine substitutions at Asp5 and Glu7 in the gp5 exonuclease domain (gp5) (32). Both complexes were prepared as previously described (32, 81). WTgp4 (82), gp4ΔC(83), gp4Δprimase (84), gp4-M1Δprimase (84), gp4-M4 (35) were also expressed and purified as previously reported. Gp4-M4 contains a C-terminal helicase mutation (E343Q) that abolishes nucleotide hydrolysis and subsequently ssDNA translocation, along with three N-terminal ZBD substitutions (P16K, D18H, N19R) that increase ssDNA binding affinity (35). The ZBD substitutions in gp4-M4 are located at the protein-DNA interface and are distal from regions proposed to contact gp5/trx.

DNA sequences encoding gp5-W1, gp5-E6, and gp5-E6exo+ were synthesized and subcloned by GenScript into pGP5-3 vectors with histidine tags and TEV cleavage sites on the N-terminus. Overproduction of proteins was carried out by transforming BL21 pLysS DE3 *E. coli* strain with appropriate plasmids. Cell were grown in 1L of LB media with 100 µg/mL ampicillin at 37°C and induced at an OD600 of 0.6 by addition of 1 mM isopropyl-1-thio-β-d-galactopyranoside (IPTG). Following induction for 4 h at 37°C, cell were harvested and resuspended in buffer A containing 20 mM potassium phosphate (pH 7.4), 10% glycerol, 50 mM β-mercaptoethanol, and 0.1 mM PMSF. Cells were combined with purified trx and lysed by sonication. Lysate was then spun at 40k rpm for 30 minutes at 4°C. The supernatant was diluted in buffer A including 40 mM imidazole and loaded to a His-Trap HP column (Cytiva) equilibrated with buffer A including 40 mM imidazole. The protein was eluted with a gradient of 50 to 600 mM imidazole, and fractions were assessed by SDS PAGE. Fractions containing gp5/trx were pooled, concentrated in 10 kDa MWCO Amicon Ultra filters (Sigma-Aldrich), and dialyzed to a storage buffer containing 50 mM ACES (pH 7.5), 50% glycerol, 1 mM EDTA, and 5 mM β-mercaptoethanol.

### DNA Ligands

Oligomers were obtained from Integrated DNA Technologies (Coralville, IA). Purity of DNA was assessed by 5’-^32^P-labeling (see **5’-^32^P-labeling**), followed by native PAGE. Sequences of all DNA ligands used in this study are presented in **SI Appendix**, **Table S1**.

### 5’-^32^P-labeling

20 µM ssDNA was mixed with 0.5 U/µL of T4 polynucleotide kinase (New England Biolabs) in a buffer containing 10 µCi [γ-^32^P] ATP, 70 mM Tris-HCl (pH 7.6) 10 mM MgCl2, and 5 mM DTT. The reaction was incubated at 37°C for 30 minutes, followed by incubation at 65°C for 20 minutes. The 5’-^32^P-labeled DNA was then purified using a Microspin G-25 Column (Cytiva).

### Fork-shaped DNA

DNA fork substrates were mixed in a 1:1.25:1 molar ratio of leading-strand, lagging-strand, and leading-strand primer, respectively, in a buffer containing 100 mM NaCl and 50 mM Tris-Cl, pH 7.5. The mixture was incubated at 95°C for 5 minutes and cooled at room temperature for several hours.

### Electrophoretic Mobility Shift Assay

Increasing concentrations of gp4-M4 (0.08-0.83 µM; hexameric concentration) or gp5/trx variants (0.1-1.0 µM) were incubated with 0.3 µM 5’-^32^P-labeled DNA substrate in a buffer containing 50 mM Tris (pH 7.5), 100 mM KCl, 5 mM DTT, 10 mM MgCl₂, and either 0.1 or 1.8 mM dTTP, as indicated. Reactions were incubated at 37 °C for 10 min, quenched by placing on ice for 5 min, and mixed with loading buffer L containing 12.5 mM Tris-HCl (pH 6.8), 8% glycerol, and 0.002% bromophenol blue. Samples were applied to 7.5% native polyacrylamide gels and run in Tris-glycine running buffer for 90 min at 130 V at 4 °C. Gels were dried and imaged using an Amersham Typhoon imager (Cytiva), and band intensities were quantified with ImageQuant TL 10.2 (Cytiva). The fraction of DNA bound was plotted as a function of protein concentration and fit to a hyperbolic equation to determine the maximal binding (B_max_) and dissociation constant (K_d_), according to:

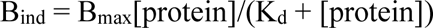

where B_ind_ is the amount of protein-DNA complex at each protein concentration, B_max_ is the maximal binding, and K_d_ is the apparent dissociation constant.

We also considered the binding of gp5/trx to its second low-affinity binding-site at the 3’-OH of double-stranded ends of the DNA fork (**SI Appendix, Fig. S5** and **S6**) and fitted binding curves with the equation reflecting a two-site binding model. This procedure had no effect on the values obtained by fitting the binding curves with a single-site model.

### Polymerization Assay

Polymerization activities of gp5/trx variants were measured on a primer-template substrate obtained from annealing a 58-nt ssDNA template strand with 20 nt ssDNA primer (5′-end labeled with ³²P). 0.3 µM primer-template was added to buffer P containing 50 mM Tris-HCl (pH 7.5), 100 mM KCl, 5 mM DTT, 10 mM MgCl2, and 0.1 mM or 1.8 mM each of dATP, dTTP, dCTP, and dGTP. Reactions were initiated by the addition of 0.3 µM gp5/trx variant (gp5/trx, gp5-E6/trx, or gp5-W1/trx) and incubated for 10 minutes at 37°C. The reaction was quenched with 20 mM EDTA and incubated on ice for 5 minutes. For denaturing gel electrophoresis, the reaction was heated to 100 °C for 30 minutes and resuspended in 80% formamide, 20 mM EDTA, and 0.02% bromophenol blue. 20 µL of each reaction were loaded to an 8M urea, 20% polyacrylamide gel and run for 16 hours at 280 V in Tris-glycine running buffer. Gels were dried and imaged using an Amersham Typhoon Biomolecular Imager (Cytiva), and band intensities were quantified with ImageQuant TL 10.2 (Cytiva). Full-length product formation was calculated as the fraction of full-length product band intensity relative to the total intensity of substrate and product bands.

### Strand displacement assay

Strand-displacement activity was measured using a DNA fork substrate obtained by annealing 43-nt and 40-nt oligonucleotides corresponding to the leading-and lagging-strands, respectively, with a 20-nt primer labeled at the 5′ end with ³²P. For experiments shown in **Fig. 5**, strand-displacement activity was assessed using DNA fork substrates containing 5′ overhangs of 0-20 nt, obtained by annealing a 37-nt leading-strand, a 20-nt 5-³²P-labeled primer, and lagging-strands of varying lengths (13-33 nt). Reactions were initiated by the addition of 0.3 µM gp5/trx variant (gp5/trx, gp5-E6/trx, or gp5-W1/trx) and either 0.1 or 1.8 mM of each dNTP to 0.3 µM DNA fork substrate in buffer P. Where indicated (**Fig. 5**), ddATP was substituted for dATP to terminate strand displacement at position N_+1_ within the parental duplex. Reactions were incubated at 37 °C for the indicated time points (30, 60, 90, and 120 s) and quenched with 20 mM EDTA. Samples were prepared for denaturing gel electrophoresis and analyzed as described in the **Polymerization assay**.

### Stop-trap assay

DNA fork substrates with varying ssDNA gaps (2-25 nt) between the fork junction and leading-strand primer-template were assembled by annealing a 40-nt ssDNA lagging strand, a 20-nt ssDNA primer (5′-end labeled with ³²P), and leading-strands of varying ssDNA lengths (35-58 nt). To assemble gp4 on the DNA fork, 0.83 µM gp4-M4 or gp4-M1Δprimase (hexameric concentration) was mixed with 0.3 µM DNA fork substrate and 0.1 mM dTTP in buffer P. The mixture was incubated for 10 min at 37 °C. DNA synthesis was initiated by adding 0.3 µM gp5/trx and 0.1 mM each of dCTP, dGTP, and ddATP. Reactions were incubated at 37°C for defined time points (30, 60, 90, 120, and 300 s) and quenched with 20 mM EDTA. Samples were prepared for denaturing gel electrophoresis and analyzed as described in the **Polymerization assay**.

### Thermal shift assay

Thermal shift assays were performed to assess the thermal stability of gp5/trx variants under different binding conditions. For baseline measurements, 1 µM gp5/trx variant (gp5/trx, gp5-E6/trx, or gp5-W1/trx) was incubated at 25 °C for 10 minutes in buffer T containing 50 mM Tris-HCl (pH 7.5), 100 mM KCl, 5 mM DTT, 10 mM MgCl₂, and 5x SYPRO Orange dye (Invitrogen). Samples were then gradually heated from 25 °C to 95 °C using a QuantStudio 3 Real-Time PCR System (Thermo Fisher Scientific), while fluorescence was continuously monitored. Melting temperatures (Tₘ) were determined by fitting the first derivative of fluorescence with respect to temperature using Protein Thermal Shift™ software (Thermo Fisher Scientific).

In assays containing gp5/trx and DNA, a DNA fork substrate with a 15-nt ssDNA gap was first prepared by annealing a 68-nt ssDNA leading-strand, a 48-nt ssDNA lagging-strand, and a 32-nt ssDNA primer. To prevent primer extension, a chain-terminating ddCTP was incorporated at the 3′ end of the primer. To determine the appropriate protein:DNA ratio, 1 µM gp5/trx was mixed with varying concentrations of DNA fork (0.5-5 µM) in buffer T with 1.8 mM dTTP (the next incoming nucleotide) and analyzed as described above. A 1:1 protein:DNA molar ratio (1 µM gp5/trx:1 µM DNA fork) produced the largest Tₘ shift, with no further increase at higher DNA concentrations, indicating saturation. This ratio was used to assess the Tₘ of complexes formed by all gp5/trx variants and DNA.

To assess leading-strand complex formation, gp4-M4 was assembled on the DNA fork prior to the addition of gp5/trx. The molar ratio of gp4-M4:DNA fork was first optimized in the same manner as described above. 1.0 µM gp4-M4 (hexameric concentration) was mixed with varying concentrations of DNA fork (0.5 - 5 µM) in buffer T with 1.8 mM dTTP and incubated at 25 °C for 10 minutes. Samples were then analyzed as described above. Analysis revealed that a 6:2 molar ratio of gp4-M4:DNA fork, corresponding to 1 µM gp4-M4 (hexameric concentration) to 2 µM DNA fork, produced maximal shift, with no additional increase in Tₘ at higher DNA concentrations. Under these conditions, 1 µM gp4-M4 (hexameric concentration) was pre-incubated with 1 µM DNA for 10 minutes at 25 °C, followed by titration of gp5/trx (1-5 µM). Mixtures were incubated for an additional 10 minutes at 25 °C to allow complex formation, and thermal shift assays were performed as described above. The optimal stoichiometry (1 µM gp5/trx:2 µM DNA:1 µM gp4-M4 hexameric concentration) was used for all subsequent experiments with different gp5/trx variants.

### Minicircle assay

The minicircle DNA substrate was prepared as described previously (59). 80 nM gp5/trx variant, 10 nM gp4 variant (hexameric concentration), and 100 nM minicircle DNA were added to a buffer containing 20 mM Tris-HCl (pH 7.5), 40 mM K-glutamate (pH 8), 5 mM DTT, 40 mM KCl, 2 mg/mL BSA, and 1.8 mM each of dATP, dCTP, dGTP, and dTTP. [α-^32^P]dGTP was present at 600 mCi/mmol. The reaction was initiated by the addition of 10 mM MgCl_2_, incubated at 37°C for a given time period (1, 5, 10, 30 min), and quenched with 20 mM EDTA. To quantify DNA synthesis, 15 µL of the reactions were spotted to DE81 filter paper and washed with ammonium formate (pH 8) three times followed by 95% EtOH. Filters were dried and [α-^32^P]dGMP incorporation was measured by scintillation counter. Background signal from buffer-only controls were subtracted from all data points. Time-course data were fit to a hyperbolic function to estimate the maximum incorporation (plateau). Processivity was calculated from the 30-minute time point by dividing total dGMP incorporation by template concentration and converting to bp based on sequence composition. In **SI Appendix**, **Fig. S11**, reaction products were denatured by heating at 95 °C for 3 min and mixed with a loading buffer containing 12.5 mM Tris-HCl (pH 6.8), 8% glycerol, and 0.002% bromophenol blue. Samples were then applied to a 0.5% alkaline agarose gel and at 50 V for 6 h in a buffer containing 50 mM NaOH and 2 mM EDTA. Gels were dried and imaged using an Amersham Typhoon Biomolecular Imager (Cytiva), and band intensities were quantified with ImageQuant TL 10.2 (Cytiva).

### Exonuclease assay

0.3 µM gp5/trx variant (gp5-exo+/trx, gp5-E6exo+/trx, or gp5-E6/trx) was mixed with 0.3 µM 54 nt ssDNA (5’-^32^P-labeled) in a buffer containing 20 mM Tris-HCl (pH 7.5), 40 mM KCl, 5 mM DTT, and 0.2 mg/mL BSA. Reactions were initiated by the addition of 10 mM MgCl₂, incubated at 37 °C, and quenched at the indicated time points (30, 60, 120, 300 s) with 20 mM EDTA. Samples were prepared for denaturing gel electrophoresis and analyzed as described in the **Polymerization assay**.

### DNA unwinding assay

dsDNA unwinding activity of gp4 variants (WT gp4, gp4Δprimase, gp4ΔC) was measured using the DNA substrate obtained by annealing a 70-nt minicircle and a 110-nt ssDNA oligonucleotide with a 5’-^32^P-label. 0.3 µM DNA substrate was mixed with 0.3 µM gp4 variant (hexameric concentration) with 0.3 µM DNA substrate in a buffer containing 50 mM Tris-HCl (pH 7.5), 100 mM KCl, 5 mM DTT, 1.6 mM dTTP, and 1.5 µM unlabeled 110-nt ssDNA to prevent re-annealing of unwound strands. Reactions were initiated with 10 mM MgCl₂ and stopped at indicated time points (1, 2, 3, 5, 10 min) with the addition of 20 mM EDTA. Samples were mixed with buffer L and loaded to a 10% nondenaturing polyacrylamide gel run in Tris-glycine running buffer for 90 minutes at 130V at 4°C. Gels were dried and imaged with an Amersham Typhoon imager (Cytiva). Bands were quantified using ImageQuant TL 10.2 (Cytiva). The fraction of DNA unwound was quantified by measuring the intensity of bands corresponding to ssDNA relative to the total DNA in each lane and plotted against time. The fraction of DNA unwound over time was fit to an exponential curve to determine the percent of maximal unwinding and unwinding rate.

### Docking of atomic coordinates into cryo-EM maps

Contour levels of all cryo-EM maps were adjusted using UCSF ChimeraX (28) such that the enclosed volume matched the theoretical molecular weight of the complex, calculated as described by Harpaz et al. (85). For the PDU maps (EMD-0391, EMD-0392, EMD-0393, EMD-0394, and EMD-0395), each representing ∼500 kDa complexes, contour levels were set to 0.024, corresponding to sigma (σ) values of 6.00, 4.80, 4.00, 6.00, and 4.80, respectively, relative to each map’s mean density. Lowering the contour below 3-3.5 σ revealed additional densities that could not be confidently interpreted and were indistinguishable from noise. The HDU map (EMD-8565) includes density for the lagging-strand DNA polymerase, which is not present in the PDU maps, and represents an ∼650 kDa complex; the contour level enclosing this complex was 0.000313, corresponding to 0.34 σ. For maps representing the PDU, atomic coordinates for helicase-ssDNA (PDB ID: 6N7V), primase (PDB ID: 1NUI), and gp5/trx-DNA fork (PDB ID: 6N7W) were fitted into the density using the “fit in map” command. All chains corresponding to DNA were removed from the coordinate files prior to fitting to enable unbiased assessment of densities corresponding to DNA, as described below. The cross-correlation coefficients (CCC) for simultaneous docking of all coordinates into the maps were 0.91 (EMDB ID: 0391), 0.73 (EMDB ID: 0392), 0.89 (EMDB ID: 0393), 0.88 (EMDB ID: 0394), and 0.86 (EMDB ID: 0395). For the map representing the HDU, improved fitting was achieved by docking individual domains of gp4 and gp5/trx, rather than entire atomic models, as described in Kulczyk at el. (3). Briefly, DNA helicase and primase (PDB IDs: 1CR0 and 1NUI) were separated into individual monomers for fitting. To fit atomic coordinates into the density representing the leading-strand DNA polymerase, PDB ID: 1T7P was divided into the following domains: exonuclease (residues 1-178), palm (179–228, 405–471, 585–698), thumb (229–256, 328–379), thioredoxin-binding loop (257–327), and fingers (472–584). To fit the lagging-strand DNA polymerase, coordinates were subdivided into palm and exonuclease (1–228, 405–471, 585–698), fingers (472–584), and thumb (229–404). Initial placement of each model was performed using the “fit in map” command, followed by refinement with the “fitmap sequence” command, in which components were iteratively fitted while subtracting the density corresponding to previously placed models. The CCC for simultaneous docking of all coordinates is 0.89. The docked HDU and PDU models were then iteratively refined using a combination of Phenix(29) and COOT1.1(30) with imposed noncrystallographic symmetry, secondary structure, and Ramachandran restraints.

To identify densities potentially corresponding to DNA in the cryo-EM maps, the atomic models were low-pass filtered using USCF ChimeraX (28) to the nominal resolution (FSC 0.143 criterion) of each map using the “molmap” command (EMD-0391: 11.2 Å, EMD-0392: 13.3 Å, EMD-0393: 11.8 Å, EMD-0394: 9.6 Å, EMD-0395: 13.8 Å; EMD-8565: 11.6 Å). The filtered maps were then contoured as described above and subtracted from their corresponding experimental maps using the “volume subtract” function. Remaining densities were fitted with chains corresponding to DNA from structures of helicase-DNA (PDB ID: 6N7V) and gp5/trx-DNA fork (PDB ID: 6N7W) complexes.

## Acknowledgments

We thank Alfredo Hernandez (Tufts University) for sharing the plasmid encoding gp4ΔC and Adam Voigt (Rutgers University) for assistance in purifying gp4ΔC and gp4-M1Δprimase. We are grateful to Maria Voigt (RCSB PDB) for help in preparing figures 1 and 10. This study was supported by the Charles and Johanna Bush Biomedical grant and a start-up grant from Rutgers University to A.W.K. A portion of this research was supported by National Institutes of Health grant R24GM154185 and performed at the Pacific Northwest Center for Cryo-EM (PNCC) with assistance from Claudia López and Marcelo de Farias using PNCC 160736 and PNCC 160223 research allocations awarded to A.W.K.

## SI Appendix Figures

**SI Fig. 1.**
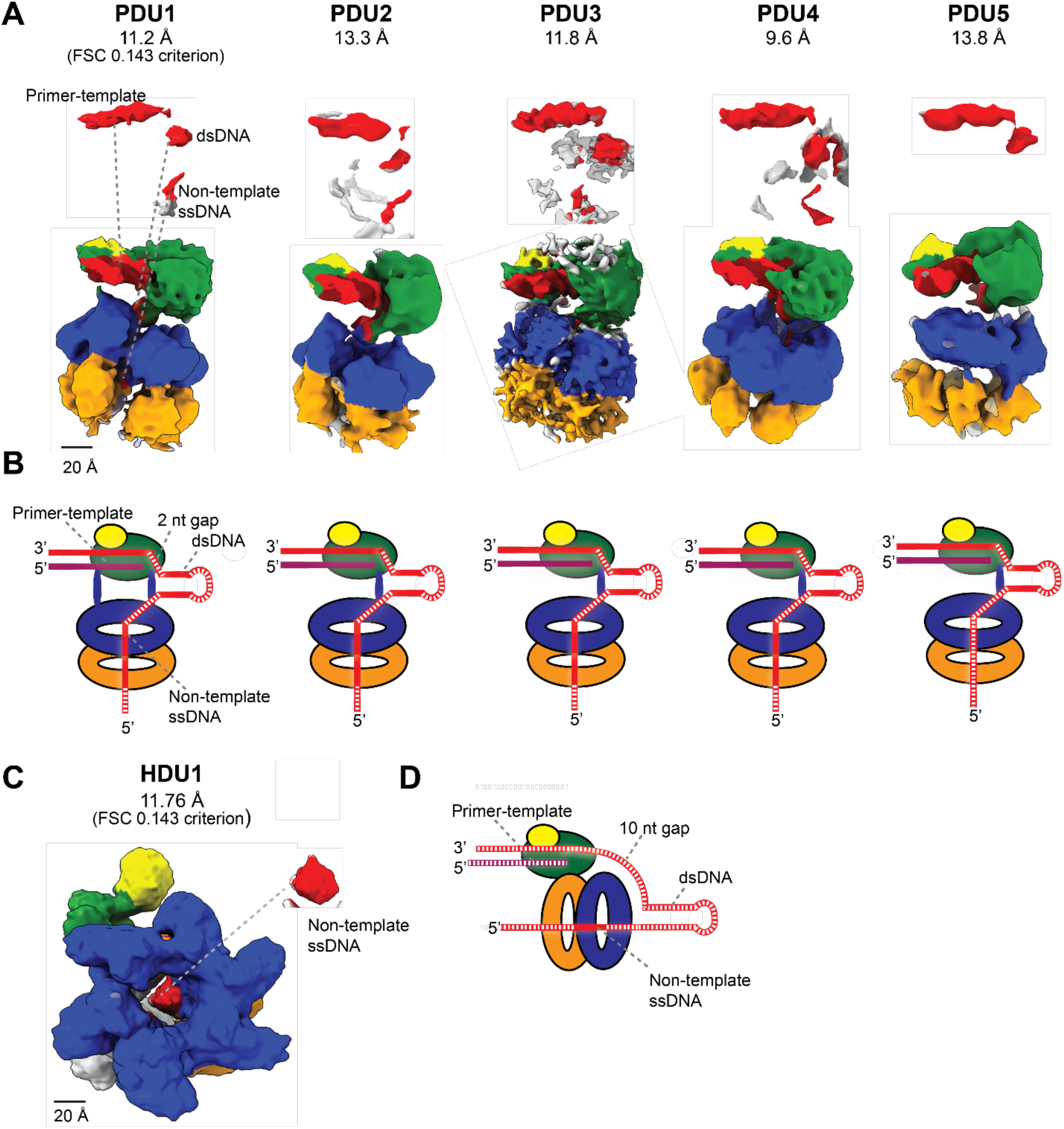
Path of the DNA fork in leading-strand complex cryo-EM reconstructions. (**A**) Cryo-EM maps representing the PDU model (PDU1-PDU5, EMDB IDs: 0391-0395), (4). Densities likely representing DNA fragments in each map are shown in red above the corresponding reconstruction (see **Materials and Methods**). DNA helicase is shown in blue, primase in orange, DNA polymerase in green, and thioredoxin in yellow. While DNA fragments are present in every map, none reveal a traceable path for the full DNA fork through both enzymes. (**B**) Cartoon representation of the DNA path through the proteins resolved in the PDU maps. Dotted lines indicate regions of DNA density that are absent or unresolved. (**C**) The cryo-EM map representing the HDU model (HDU1, EMDB ID: 8565) did not directly resolve the DNA fork substrate used in the sample, but density (red) in the central channel of the hexameric helicase may correspond to ssDNA (3). (**D**) Cartoon representation of the DNA path through the proteins resolved in the HDU map.

**SI Fig. 2.**
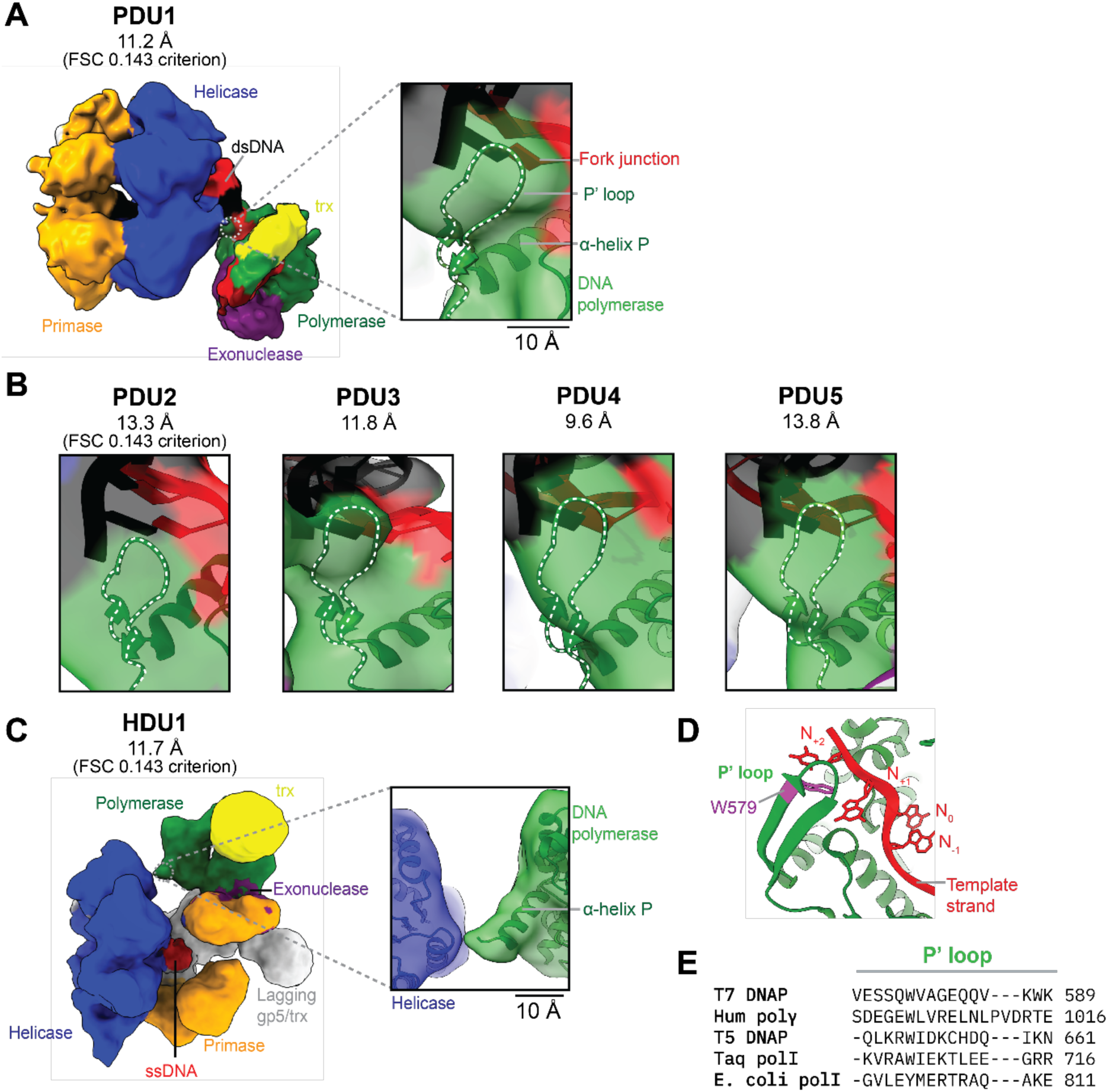
The P′ loop of DNA polymerase is not unambiguously resolved in leading-strand complex cryo-EM maps. (**A**) Cryo-EM reconstruction representing the PDU model (PDU1, EMDB ID: 0391). The inset highlights the region corresponding to the fork junction and expected position of the P′ loop, fitted with gp5-DNA fork coordinates (PDB ID: 6N7W), as described in **Materials and Methods**. (**B**) Corresponding region shown in cryo-EM maps of PDU2-PDU5 (EMDB IDs: 0392-0395), illustrating the absence of specific density for the P′ loop across all reconstructions. (**C**) The HDU1 cryo-EM reconstruction (EMDB ID: 8565) does not contain density for the P’ loop. The map is fitted with atomic coordinates for gp5/trx (PDB ID:1T7P) and helicase (PDB ID: 1CR0). (**D**) Crystal structure of the gp5/trx-primer-template complex (PDB ID:2AJQ) shows W579 (magenta) of the P′ loop base stacking with the template base two nucleotides downstream (N_+2_) from the nucleotide positioned in the polymerization active site (N_0_). (**E**) Amino acid sequence alignment of selected DNA polymerases showing the P′ loop (residues Glu575-Gln585). Included polymerases: T7 DNA polymerase (T7 DNAP), Homo sapiens DNA polymerase γ (Hum polγ), T5 DNA polymerase (T5 DNAP), Thermus aquaticus DNA polymerase I (Taq polI), and Escherichia coli DNA polymerase I (E. coli polI). Sequences were aligned using Clustal Omega 1.83 (86).

**SI Fig. 3.**
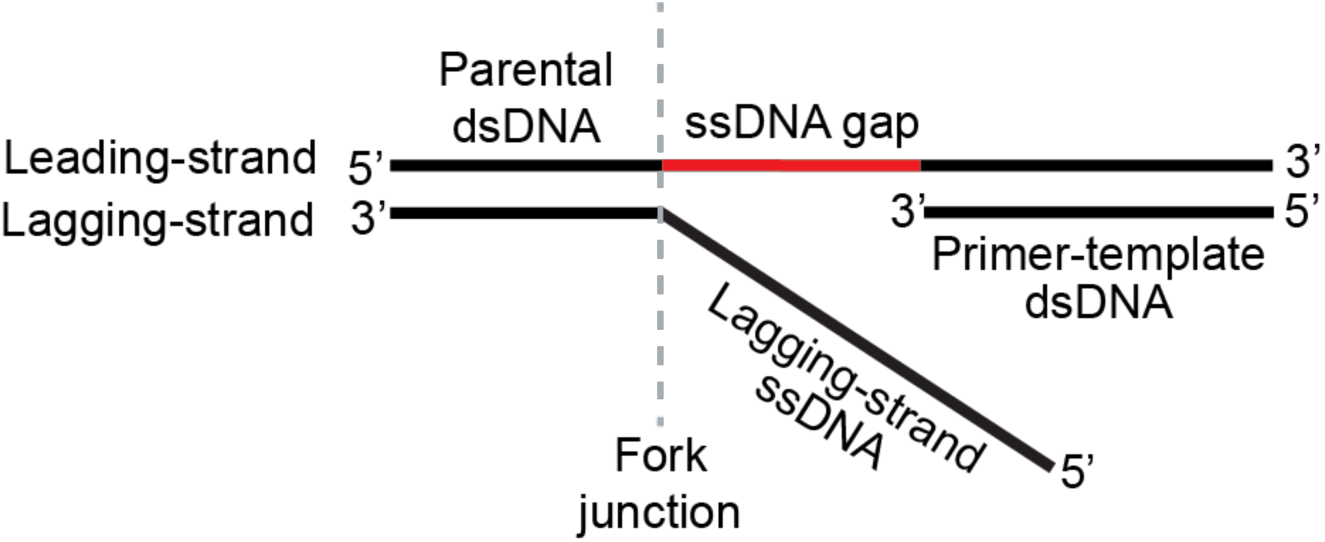
Architecture of simplified replication fork for leading-strand synthesis. The DNA fork includes parental dsDNA, a leading-strand dsDNA primer-template, a ssDNA gap between the primer-template and fork junction, and a lagging-strand ssDNA overhang. The primer-template serves as a binding site for DNA polymerase and provides a 3′-OH required for DNA polymerase to continuously copy the leading-strand. The lagging-strand 5’-overhang provides a binding site for DNA primase-helicase.

**SI Fig. 4.**
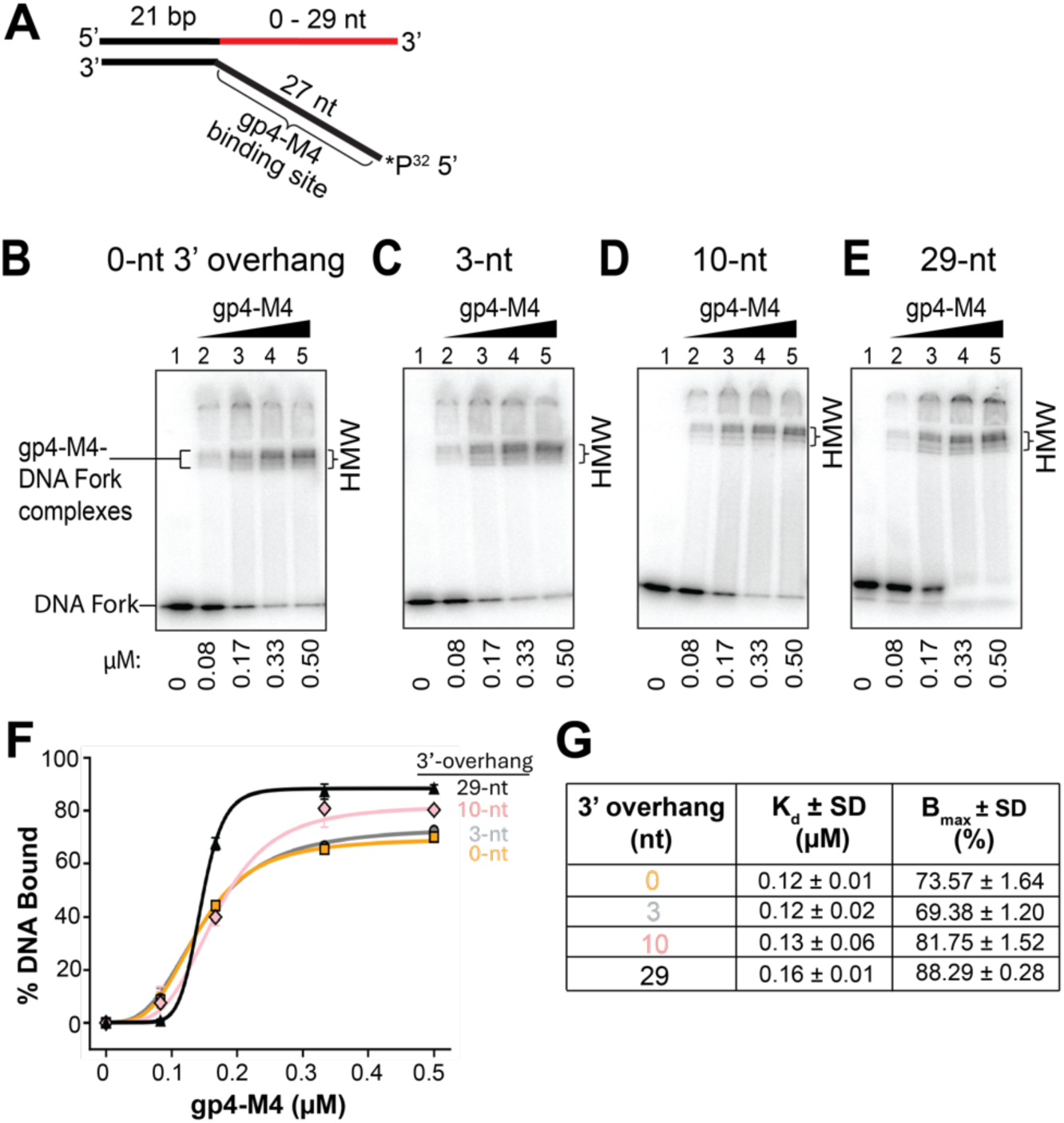
Characterization of gp4-M4 binding to DNA fork substrates with varying 3’-overhang lengths. (**A**) DNA forks with different 3’-overhang lengths (0-29 nt) were assembled by annealing leading strands of varying lengths (21-50 nt) to a 48-nt lagging-strand (5’-end labeled with ³²P). DNA forks (0.3 µM) were incubated with increasing amounts of gp4-M4 (0.083-0.5 µM, hexameric concentration) in the presence of 0.1 mM dTTP at 37 °C for 10 minutes. (**B-E**) Representative EMSA gels revealing minimal protein stacking in the well. High-molecular-weight (HMW) bands representing gp4-DNA complexes are indicated. Multiple bands likely represent different loading conformations that have been observed previously (38). Faint bands migrating below the unbound DNA fork on the gel represent excess unannealed 5’-³²P-labeled lagging-strand. (**F**) Quantification of DNA binding. The fraction of gp4-M4 bound to DNA was quantified by measuring the intensity of all HMW bands relative to total DNA in each lane and plotted against protein concentration. Error bars represent standard deviation of three independent experiments. (**G**) Binding parameters for gp4-M4 derived from curve fits, including maximum binding (B_max_) and dissociation constants (K_d_).

**SI Fig. 5.**
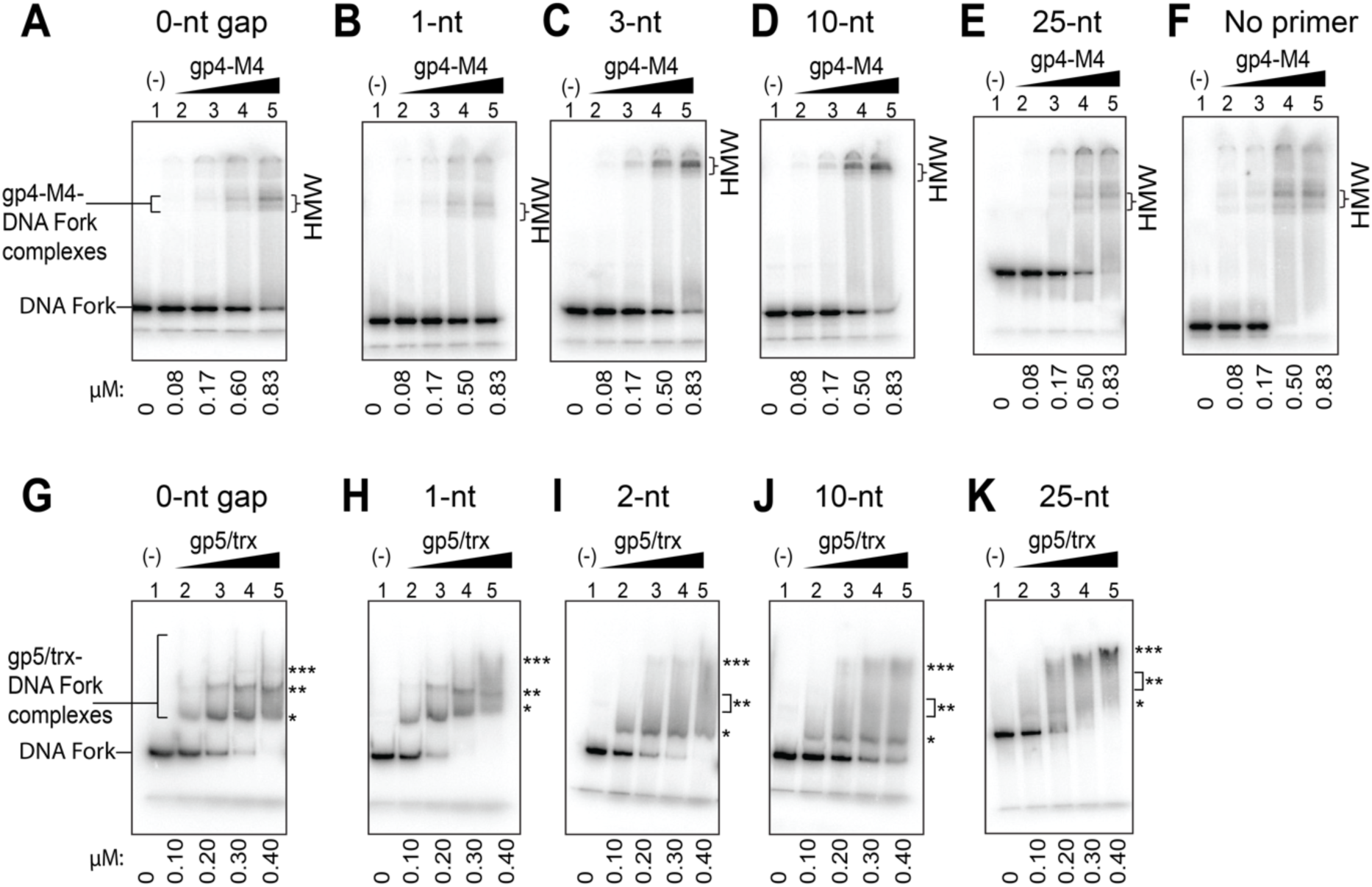
Characterization of DNA binding affinity of gp4-M4 and gp5/trx for DNA forks with varying ssDNA gap sizes. DNA fork substrates containing ssDNA gaps of 0-25 nt between the fork junction and the primer-template were assembled as described in Fig. 4. DNA ligands (0.3 µM) were incubated with increasing concentrations of gp4-M4 (0.08-0.83 µM, hexameric concentration) or gp5/trx (0.1-0.4 µM) in the presence of 0.1 mM dTTP at 37 °C for 10 min. (**A-F**) Representative EMSA gels of gp4-M4. High-molecular-weight (HMW) bands representing gp4-DNA complexes are indicated. Multiple bands likely represent different loading conformations that have been observed previously (38). Faint bands migrating below the unbound DNA fork on the gel represent excess unannealed 5’-³²P-labeled primer. (**G-K**) Representative EMSA gels of gp5/trx. Multiple bands corresponding to gp5/trx-DNA complexes are consistent with data reported previously (21, 47), and have been proposed to represent gp5/trx binding to other structural features of DNA. *Represents specific gp5/trx binding to its high-affinity 3’-end of the primer-template. **Represents the above complex with an additional copy of gp5/trx likely binding to the 3’-OH at one of the double-stranded ends of the DNA fork with a higher off-rate, as indicated by band smearing. ***Represents fork DNA with at least three molecules of gp5/trx binding, one to its high-affinity site at the 3’-end of the primer-template and additional gp5/trx molecules binding to 3’-OH groups at the two dsDNA ends. See **SI** Fig. 3.

**SI Fig. 6.**
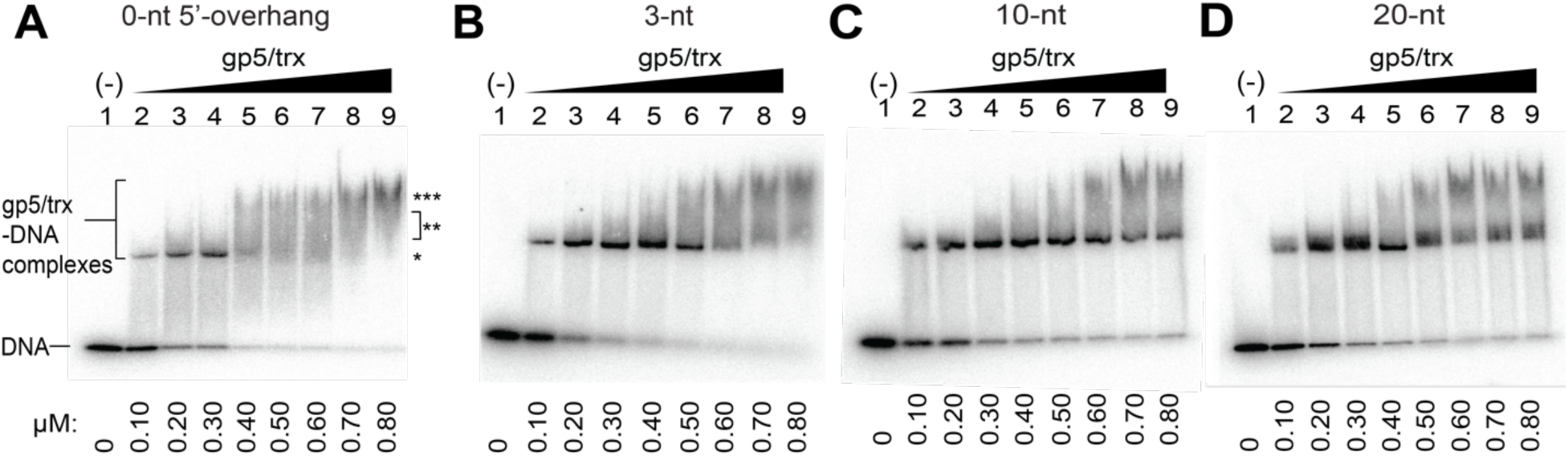
EMSA analysis of gp5/trx-stalled complexes on DNA forks with varying 5′-overhang lengths. DNA forks containing 0-, 3-, 10-, or 20-nt 5′-overhangs were assembled as described in Fig. 5A. Strand-displacement reactions were initiated by mixing 0.3 µM DNA fork with increasing amounts of gp5/trx (0.1-0.8 µM) and 0.1 mM each of dCTP, dTTP, dGTP, and chain-terminating ddATP to stop strand-displacement 2 bp inside the parental duplex. Reactions were incubated at 37 °C for 2 minutes and resolved on native polyacrylamide gels. (**A-D**) Representative EMSA gels. Multiple bands corresponding to gp5/trx-DNA complexes observed at high protein concentrations likely reflect the ability of gp5/trx to recognize different structural features of DNA, consistent with **SI** Fig. 5G**-K** and previous data (21, 47). *Represents gp5/trx binding to its high-affinity 3’-end of the primer-template. **,***Represent the above complex with additional copies of gp5/trx likely binding to the two 3’ OH groups at double-stranded ends of the DNA fork.

**SI Fig. 7.**
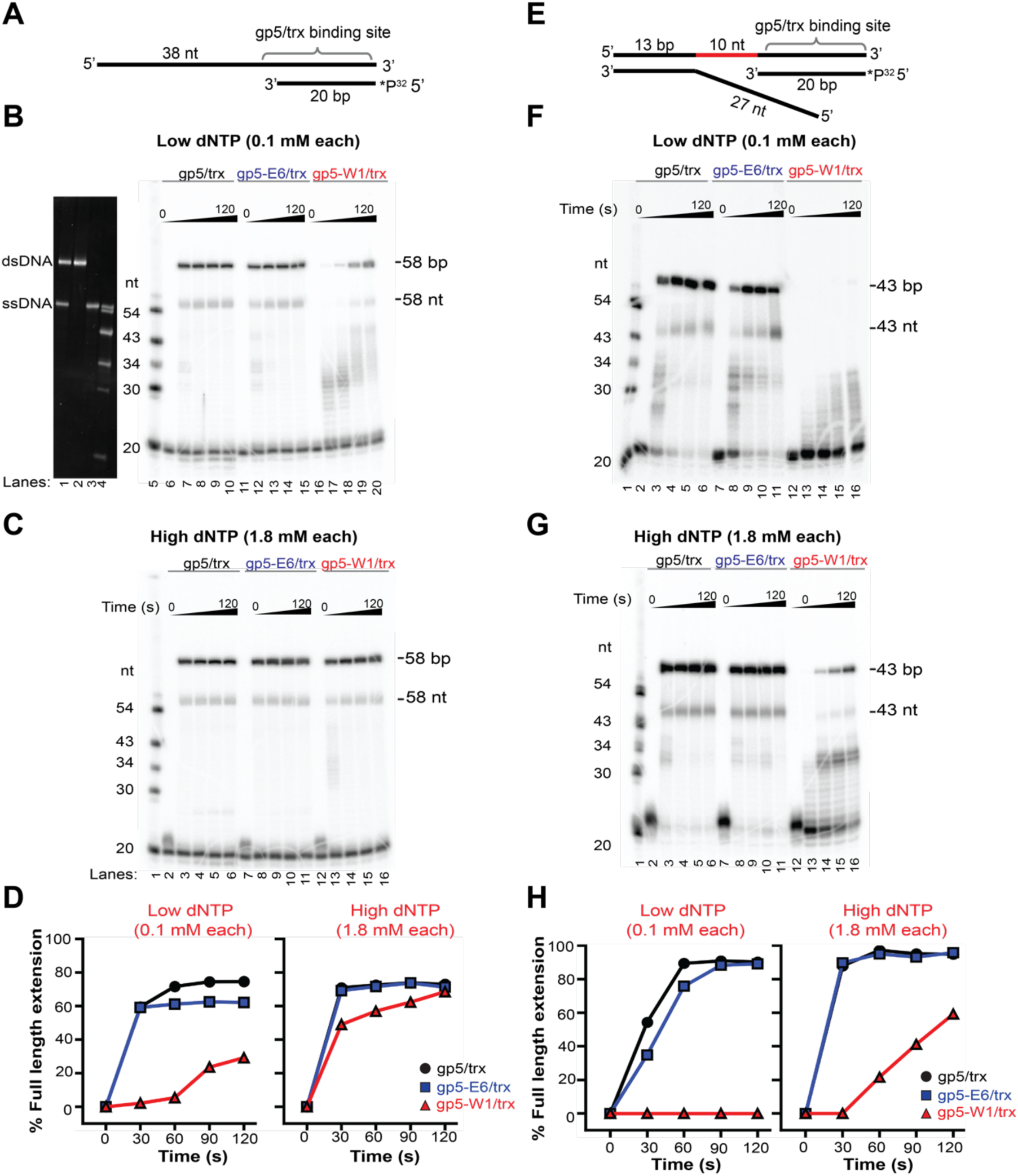
Polymerization and strand-displacement activities of gp5/trx variants at low and high dNTP concentrations. (**A**) Polymerization activities of gp5/trx variants were measured on a primer-template substrate obtained from annealing a 58-nt template strand with 20-nt primer (5′-end labeled with ³²P). Reactions were initiated by mixing equimolar amounts of each gp5/trx variant and DNA in the presence of 0.1 mM or 1.8 mM of each dNTP. Reactions were quenched at indicated time points (30, 60, 90, 120 s) with 20 mM EDTA. (**B, C**) Representative denaturing 8 M urea, 20% polyacrylamide gels showing polymerization activity of gp5/trx (lanes 7-10), gp5-E6/trx (lanes 12-15) and gp5-W1/trx (lanes 17-20) at the low (**B**) and high (**C**) dNTP concentrations. Despite denaturing conditions, a substantial fraction of reaction products corresponding to fully extended 58-nt DNA products remained double-stranded. Control DNA fragments in **B** were visualized by SYBR gold staining and include a mixture of 58-bp dsDNA and 58-nt ssDNA (lane 1), 58-bp dsDNA alone (lane 2), 58-nt ssDNA alone (lane 3), and a ladder (lane 4), confirming the identity of the fully extended 58-nt products. (**D**) Quantification of full-length product formation at indicated timepoints, expressed as the ratio of the full-length product band to the total of all substrate and product bands. (**E-H**) Strand-displacement activities of gp5/trx variants were measured using a DNA fork substrate obtained by annealing a 43-nt and 40-nt ssDNA corresponding to leading- and lagging-strands, respectively, and a 20-nt primer (5′-end labeled with ³²P). Reactions were carried out as described in **A**. (**F, G**) Representative denaturing 8 M urea, 20% polyacrylamide gels showing strand-displacement activity of gp5/trx (lanes 3-6), gp5-E6/trx (lanes 8-11) and gp5-W1/trx (lanes 13-16) at the low (**F**) and high (**G**) dNTP concentrations. (**H**) Quantification of full-length product formation at indicated timepoints.

**SI Fig. 8.**
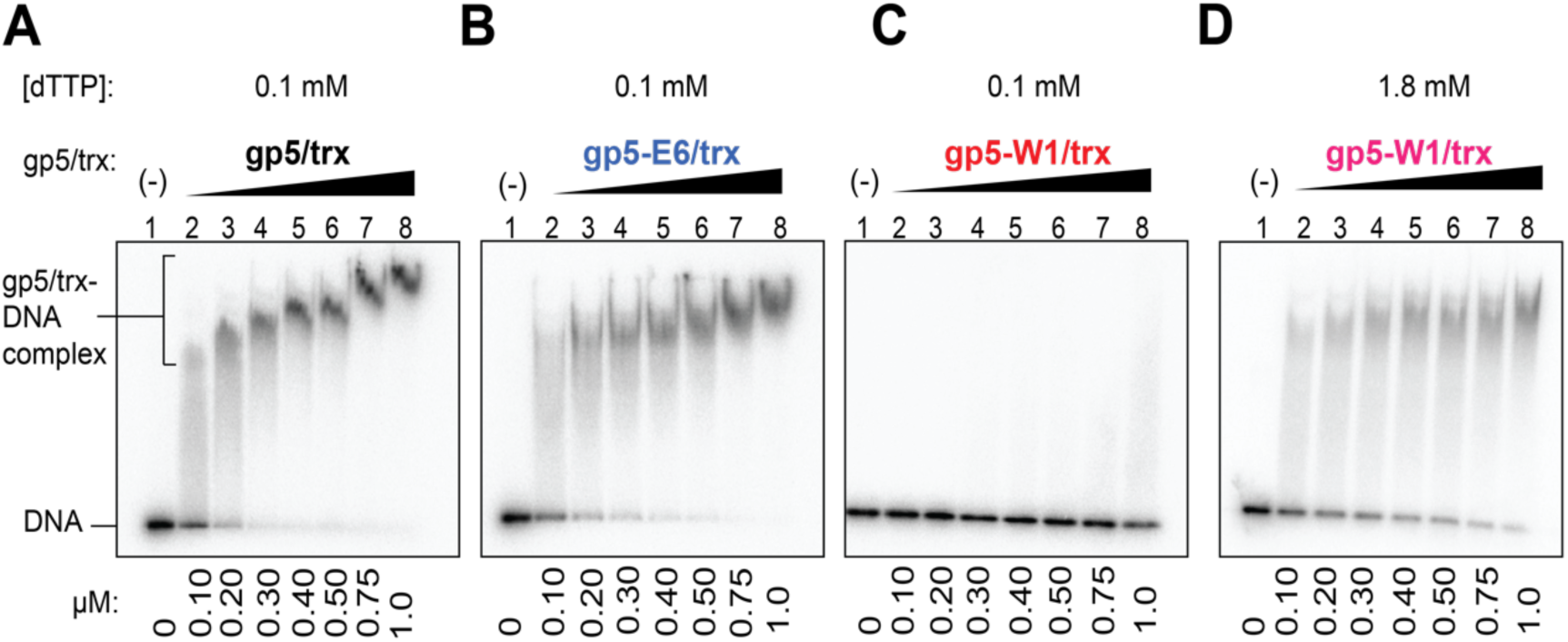
Characterization of DNA binding affinity of gp5/trx variants. EMSA was used to assess binding of gp5/trx variants to a primer-template obtained by annealing a 58-nt template strand to a 20-nt primer (5′-³²P-labeled). The primer contains a 3′-dideoxycytidine (ddC) to prevent extension. DNA (0.3 µM) was incubated with increasing amounts of gp5 variants (0.1-1.0 µM) in the presence of dTTP (0.1 or 1.8 mM) at 37 °C for 10 min. (**A-D**) Representative EMSA gels for gp5/trx (**A**), gp5-E6/trx (**B**), and gp5-W1/trx at 0.1 mM (**C**) and 1.8 mM (**D**) dTTP.

**SI Fig. 9.**
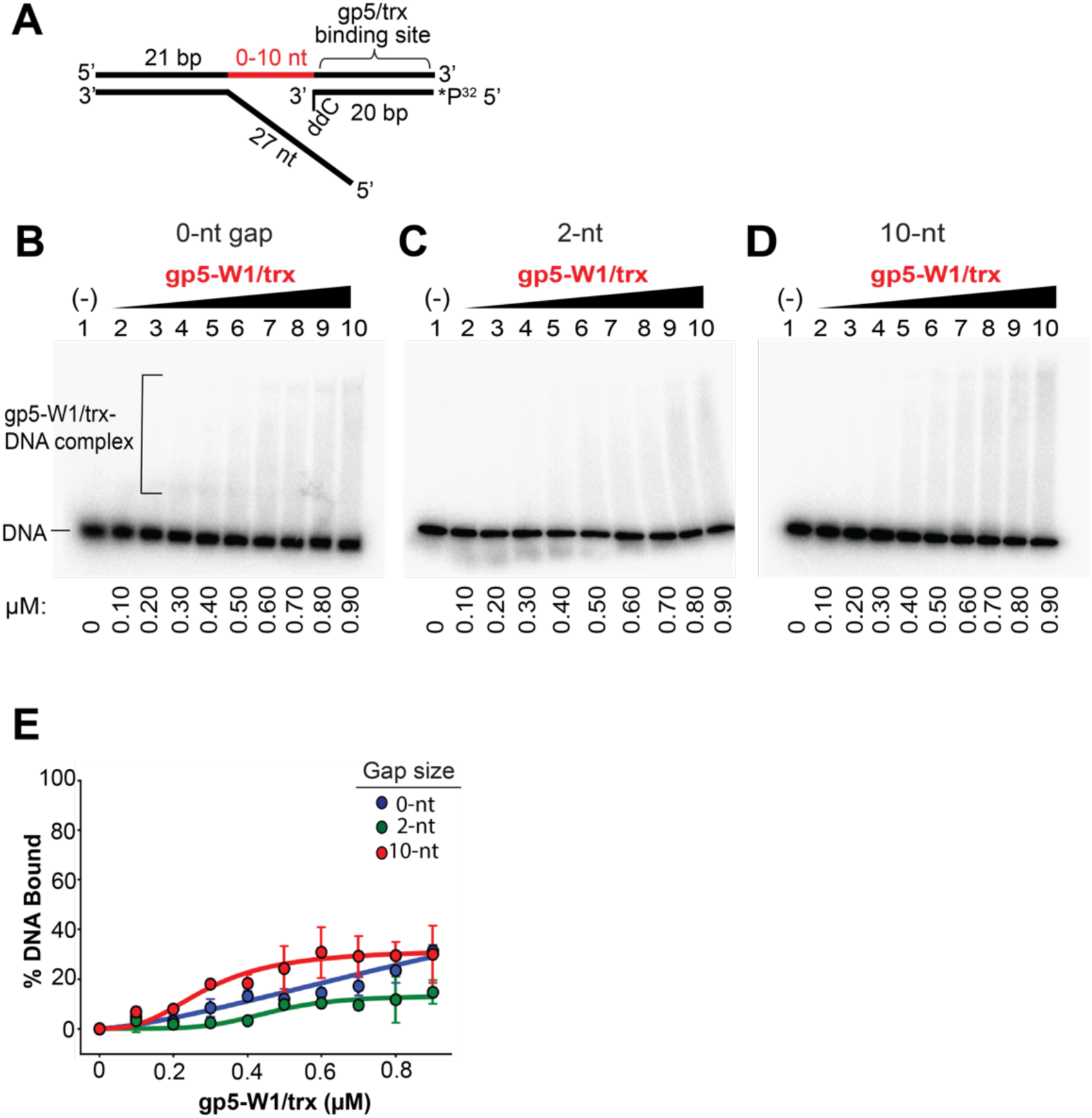
Characterization of gp5-W1/trx binding to DNA fork substrates with different size ssDNA gaps. (**A**) DNA fork substrates containing ssDNA gaps of 0-, 2-, and 10-nt ssDNA gaps between the fork junction and the primer-template were assembled as described in Fig. 4. DNA ligands (0.3 µM) were incubated with increasing amounts of gp5-W1/trx (0.1-0.9 µM) in the presence of 1.8 mM of the next incoming dNTP specified by the template strand, at 37 °C for 10 min. (**B-D**) Representative EMSA gels for each substrate. (**E**) Fraction of DNA bound plotted as a function of gp5-W1/trx concentration. Error bars represent the standard deviation of two independent experiments.

**SI Fig. 10.**
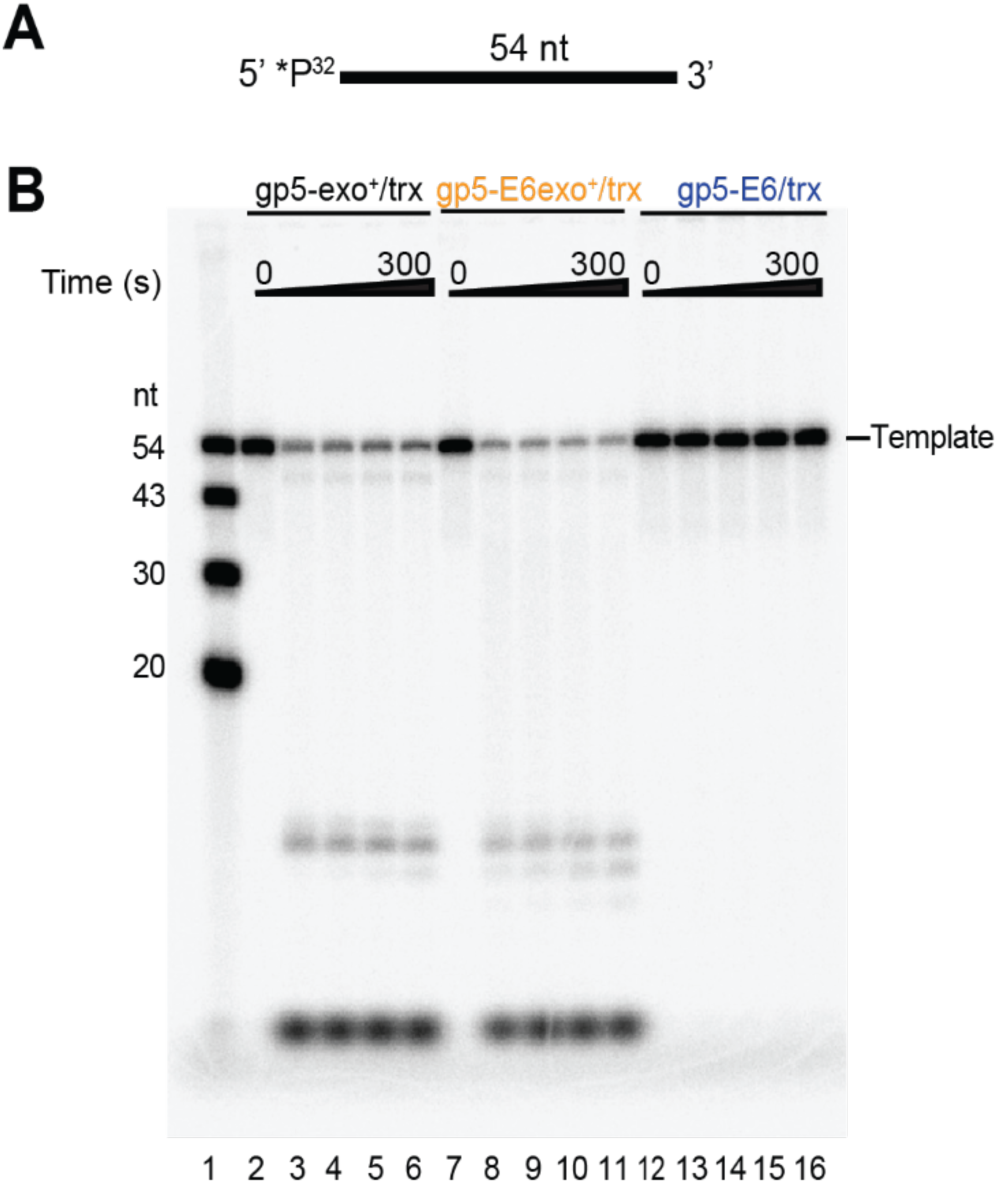
Exonuclease activity of gp5-E6/trx is unaffected by mutations in the acidic patch. (**A**) Equimolar amounts of each gp5/trx variant were mixed with a 54-nt ssDNA template (5’-^32^P-labeled). Reactions were initiated by the addition of 20 mM MgCl2 and quenched at indicated time points (30, 60, 120, and 300 s) with 20 mM EDTA. (**B**) Representative denaturing polyacrylamide gel comparing exonuclease activity of wild-type gp5/trx (gp5-exo⁺/trx; lanes 3-6), exonuclease-proficient gp5-E6/trx (gp5-E6-exo⁺/trx; lanes 8-11), and exonuclease-deficient gp5-E6/trx (gp5-E6/trx; lanes 13-16). gp5-E6-exo⁺/trx exhibits exonuclease activity comparable to gp5-exo⁺/trx, whereas no exonuclease activity is observed for the exonuclease-deficient gp5-E6/trx.

**SI Fig. 11.**
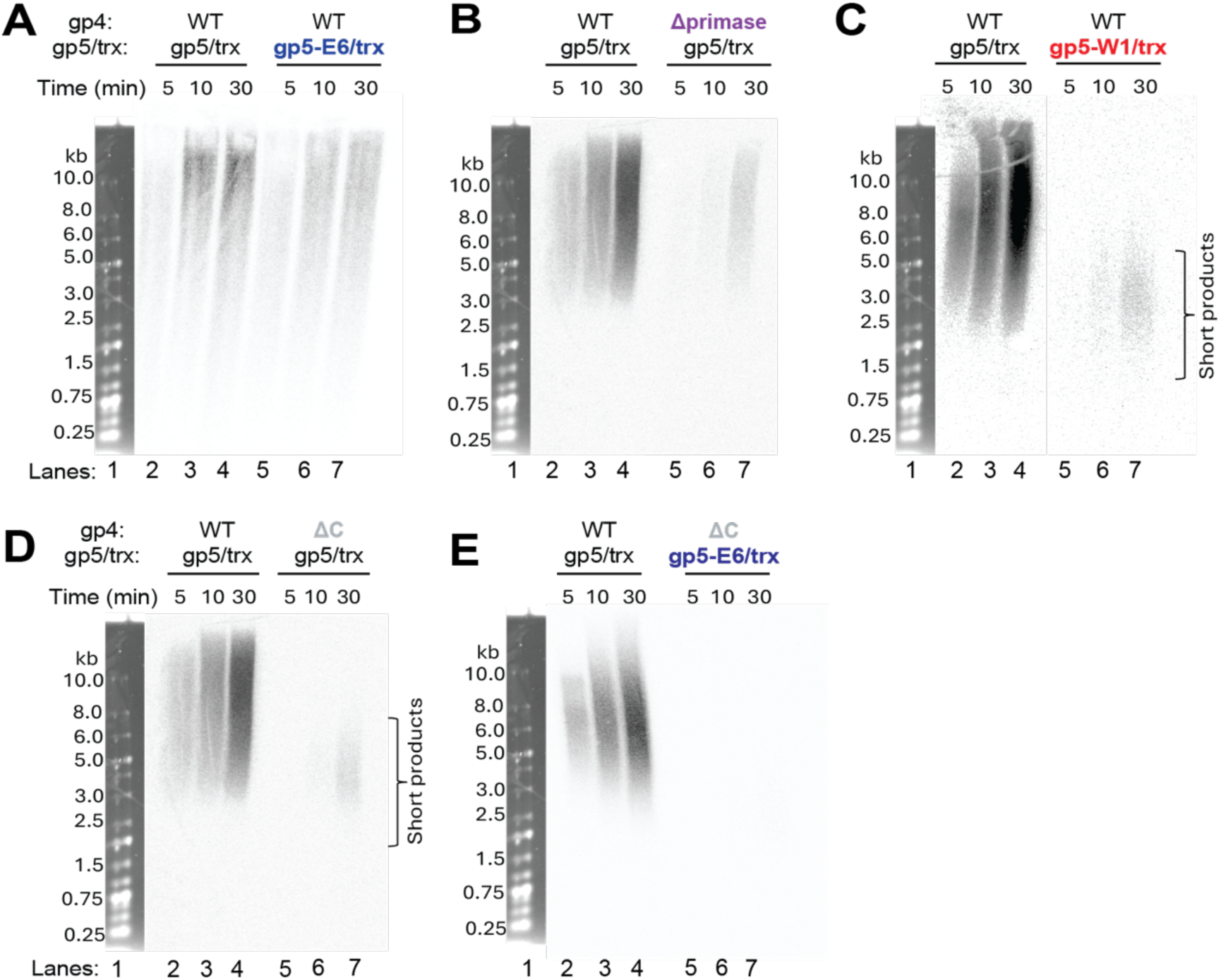
Processivity of gp5/trx and gp4 variants on minicircle DNA. (**A-E**) Reactions containing 80 nM gp5/trx variant, 10 nM gp4 variant (hexameric concentration), and 100 nM minicircle DNA were carried out in the presence of 1.8 mM each dNTP, with [α-³²P]dGTP (600 mCi/mmol) included for monitoring DNA synthesis. Reactions were initiated by the addition of 10 mM MgCl₂ and incubated at 37 °C for indicated time points (5, 10, 30 min) before quenching with 20 mM EDTA. Reaction products were denatured at 95 °C and analyzed by 0.5% alkaline agarose gel electrophoresis. Gels are displayed at exposures optimized for visualization of product bands.

**SI Fig. 12.**
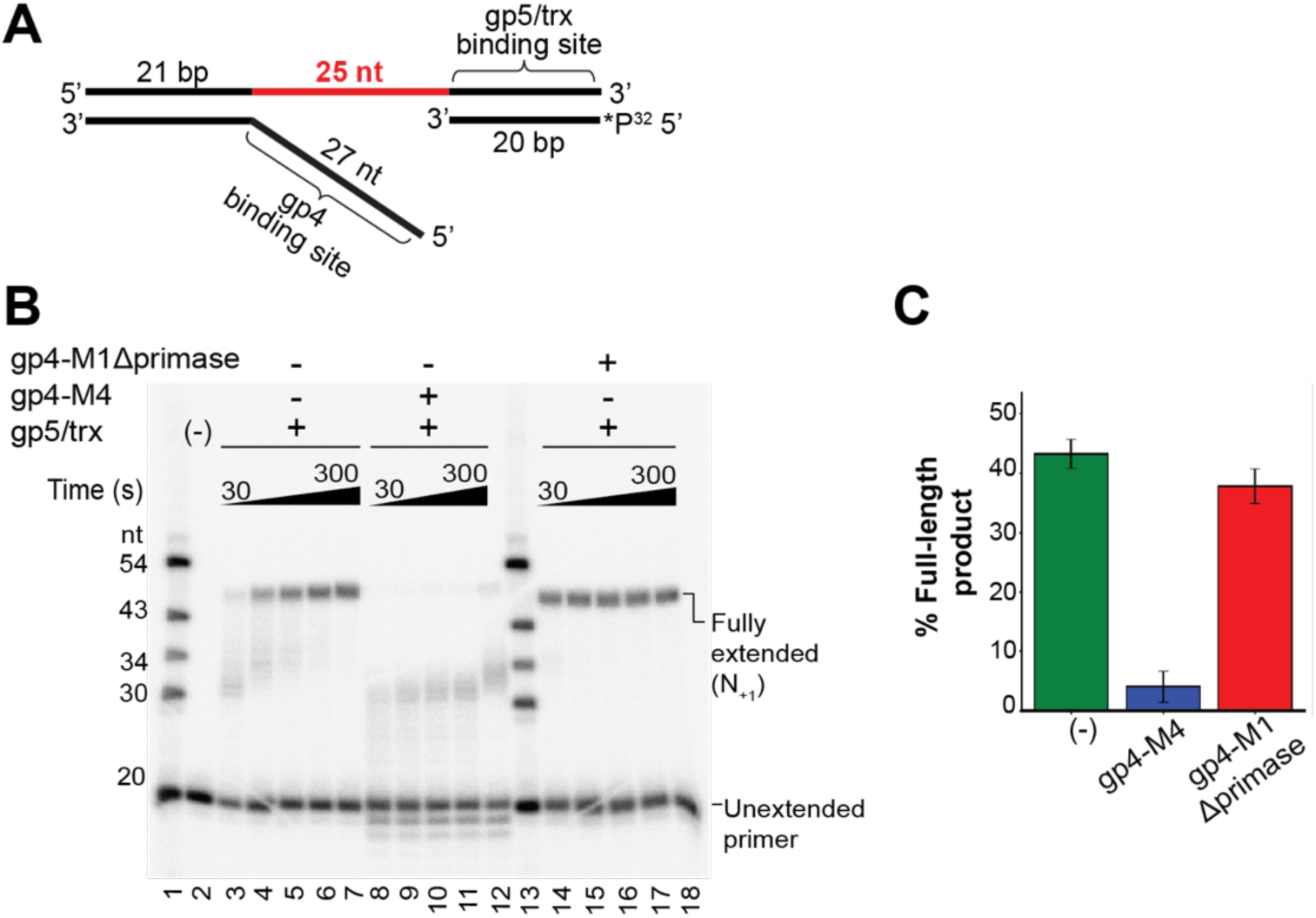
gp4-M1Δprimase does not stop DNA synthesis by DNA polymerase in the stop-trap assay. (**A**) gp4-M1Δprimase was evaluated in the stop-trap assay with gp5/trx using a DNA fork substrate containing a 25-nt gap between the primer 3’-end and the fork junction. The DNA fork was assembled as described in Fig. 6. In this reaction, 0.3 µM DNA fork was first incubated with 0.83 µM gp4-M1Δprimase or gp4-M4 (hexameric concentration) in the presence of 0.1 mM dTTP at 37 °C for 10 minutes. DNA synthesis was initiated by the addition of 0.3 µM gp5/trx together with 0.1 mM dCTP, dGTP, and chain-terminating ddATP to stop strand-displacement 2 bp inside the parental duplex. Reactions were stopped at indicated time points (30, 60, 90, 120, 300 s) with the addition of 20 mM EDTA. (**B**) The products were resolved on 8M, 20% polyacrylamide gels. Oligonucleotides of known length labeled at the 5’-end with ^32^P were included in lanes 1 and 13 to measure the size of reaction products. (**C**) Quantification of full-length product formation at 300 s was calculated as the ratio of the intensity of the full-length product band to the total intensity of all substrate and product bands. Error bars represent standard deviation of three independent experiments.

**SI Fig. 13.**
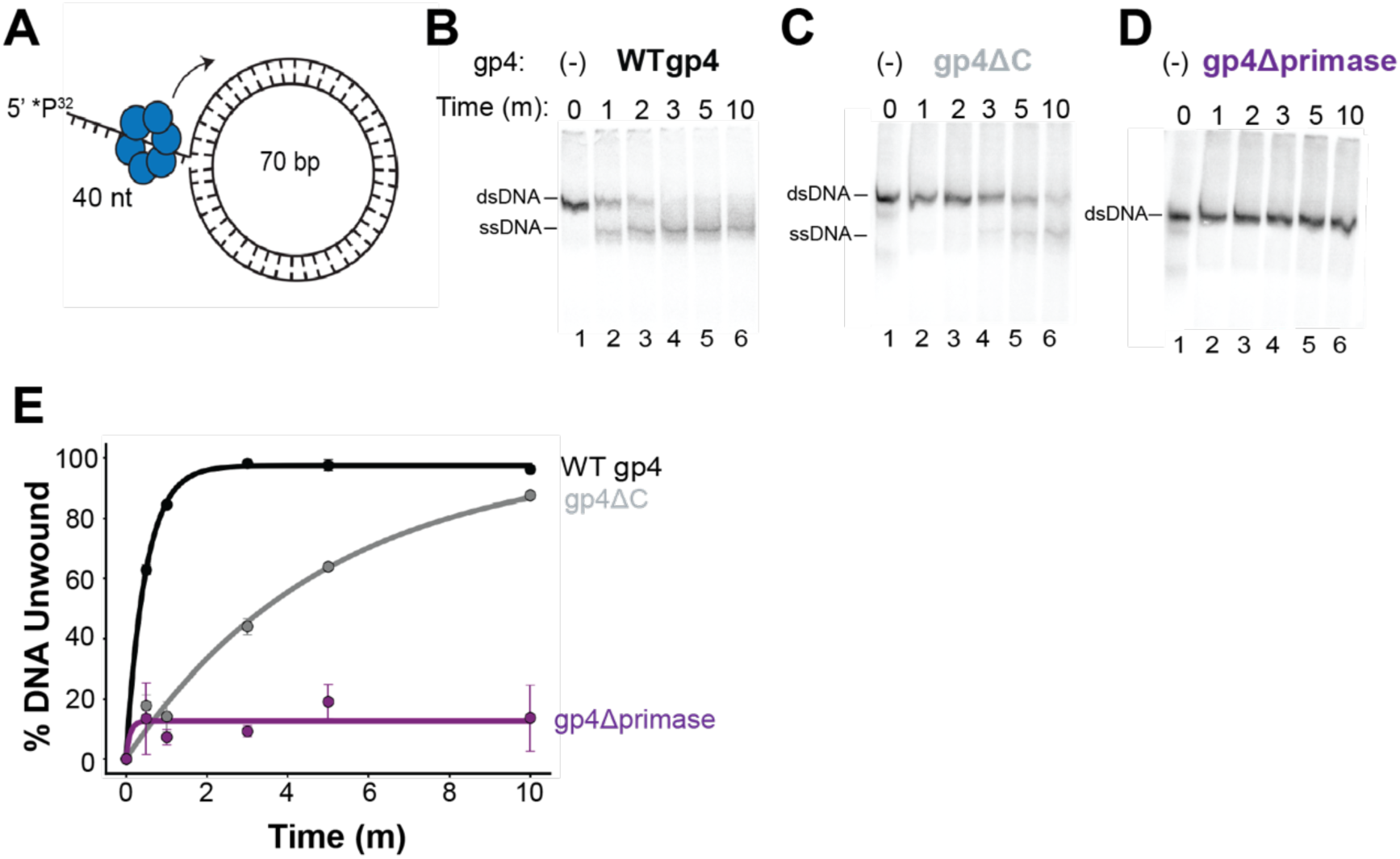
Characterization of unwinding activity of gp4 variants. (A) The DNA unwinding activity of WTgp4, gp4ΔC, and gp4Δprimase were assessed using the DNA substrate obtained upon by annealing a 70-nt ssDNA minicircle and a 110-nt ssDNA oligonucleotide with a 5’ ^32^P-label. Reactions contained 0.3 µM gp4 variant (hexameric concentration), 0.3 µM minicircle DNA, 1 mM dTTP, and 1.5 µM unlabeled 110-nt ssDNA to act as a “trap” preventing re-annealing of the unwound strands. Reactions were initiated with the addition of 10 mM MgCl_2_ and stopped at indicated time points (1, 2, 3, 5, 10 min) with the addition of 20 mM EDTA (B-D). Unwinding was assessed by native PAGE separation of 5′ ³²P-labeled ssDNA from dsDNA, for (**B**) WT gp4, (**C**) gp4ΔC, and (**D**) gp4Δprimase. (**E**) The percent of DNA unwound was quantified by measuring the intensity of bands corresponding to ssDNA relative to the total DNA in each lane and plotted as a function of time. Data points represent the mean of three independent experiments; error bars indicate standard deviation.

---

**Table 1:**
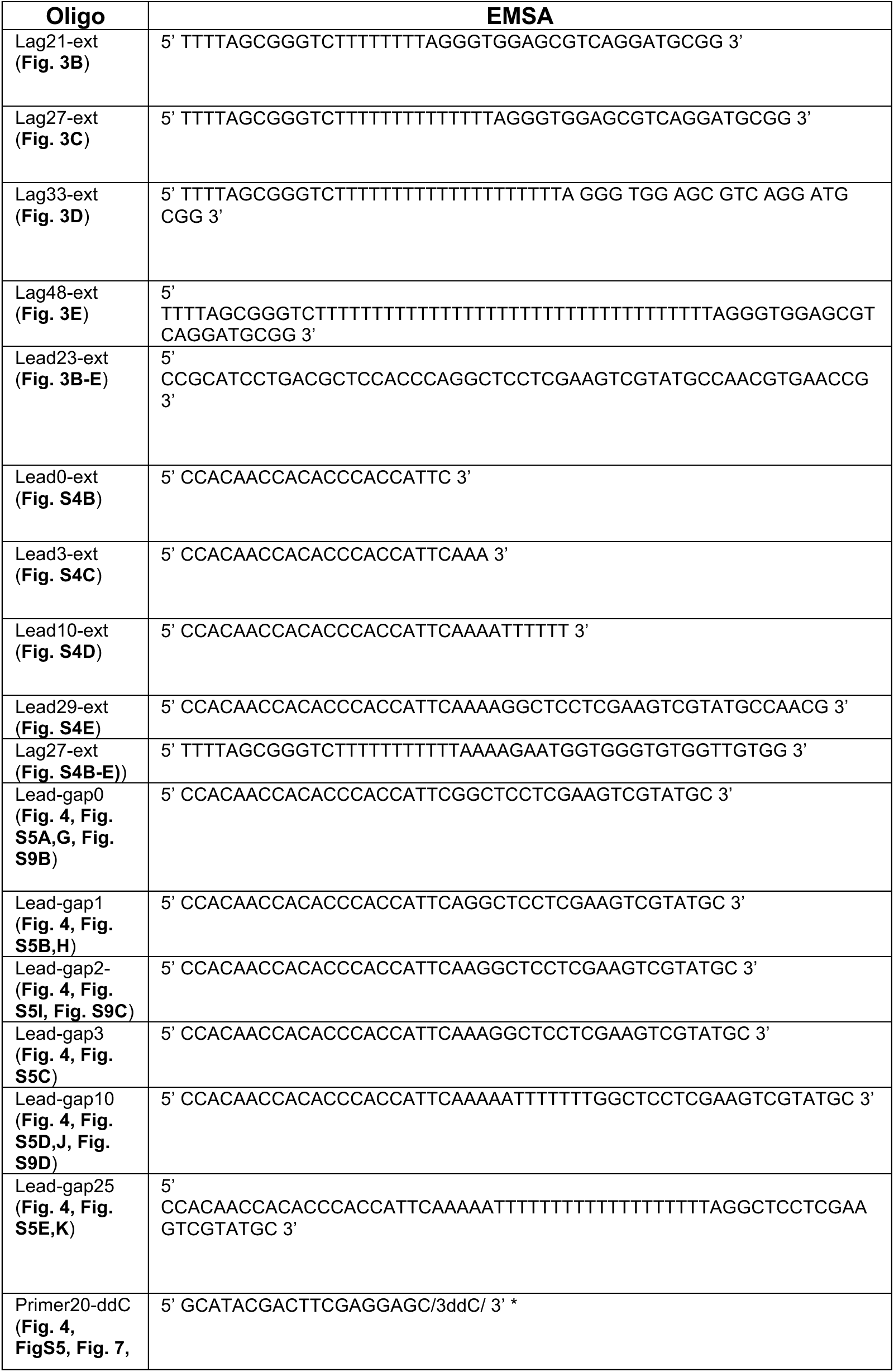
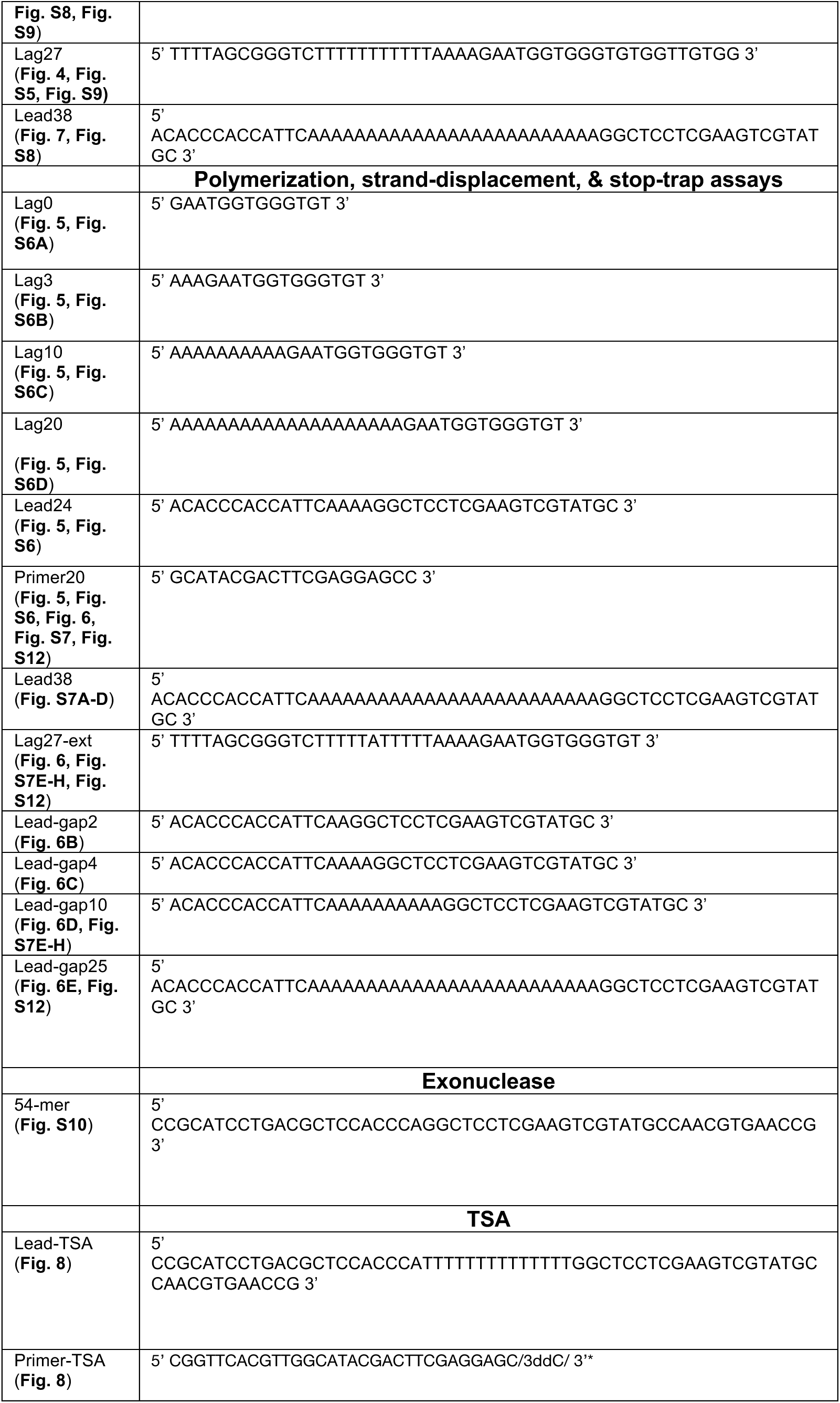
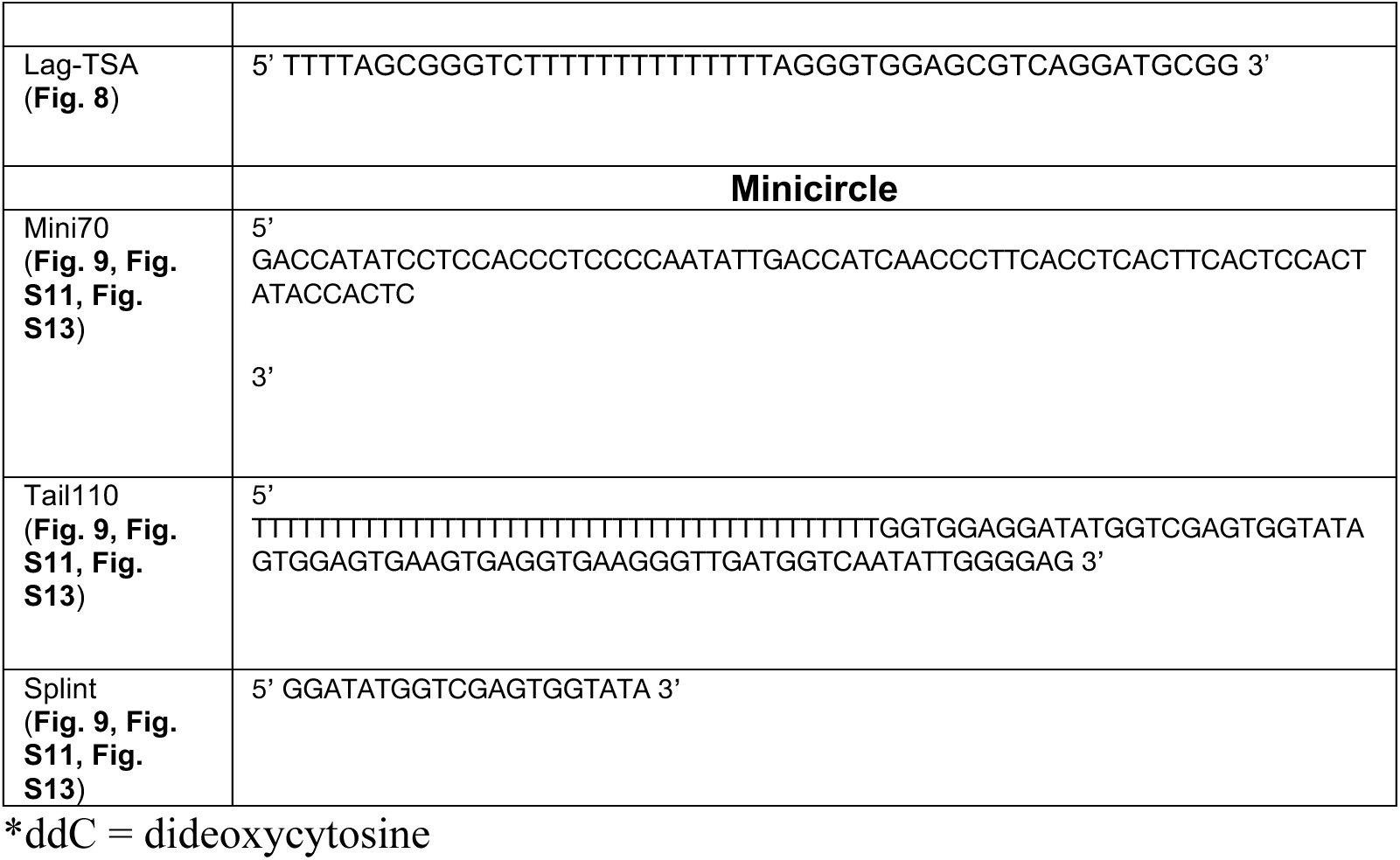
Oligomers used in EMSA, polymerization, strand-displacement, stop-trap, exonuclease, thermal shift, and rolling circle assays.

## Movies

**Movie S1** (separate file). The movie shows cryo-EM maps of the T7 replisome sub-assemblies that support PDU (EMDB IDs: 0391-0395), (4) or HDU (EMDB ID: 8565), (3) models of leading-strand replication.

